# Machine learning of molecular dynamics simulations provides insights into modulation of viral capsid assembly

**DOI:** 10.1101/2025.02.07.637202

**Authors:** Anna Pavlova, Zixing Fan, Diane L. Lynch, James C. Gumbart

## Abstract

An effective approach in the development of novel antivirals is to target the assembly of viral capsids using capsid assembly modulators (CAMs). CAMs targeting hepatitis B virus (HBV) have two major modes of function: they can either accelerate nucleocapsid assembly, retaining its structure, or misdirect it into non-capsid-like particles. Previous molecular dynamics (MD) simulations of early capsid-assembly intermediates showed differences in protein conformations for apo and bound states. Here, we have developed and tested several classification machine learning (ML) models to better distinguish between apo-tetramer intermediates and those bound to accelerating or misdirecting CAMs. Models based on tertiary structural properties of the Cp149 tetramers and their inter-dimer orientation, as well as models based on direct and inverse contact distances between protein residues, were tested. All models distinguished the apo states and the two CAM-bound states with high accuracy. Furthermore, tertiary structure models and residue-distance models highlighted different tetramer regions as important for classification. Both models can be used to better understand structural transitions that govern the assembly of nucleocapsids and to assist the development of more potent CAMs. Finally, we demonstrate the utility of classification ML methods in comparing MD trajectories and describe our ML approaches, which can be extended to other systems of interest.

**TOC Graphic:** 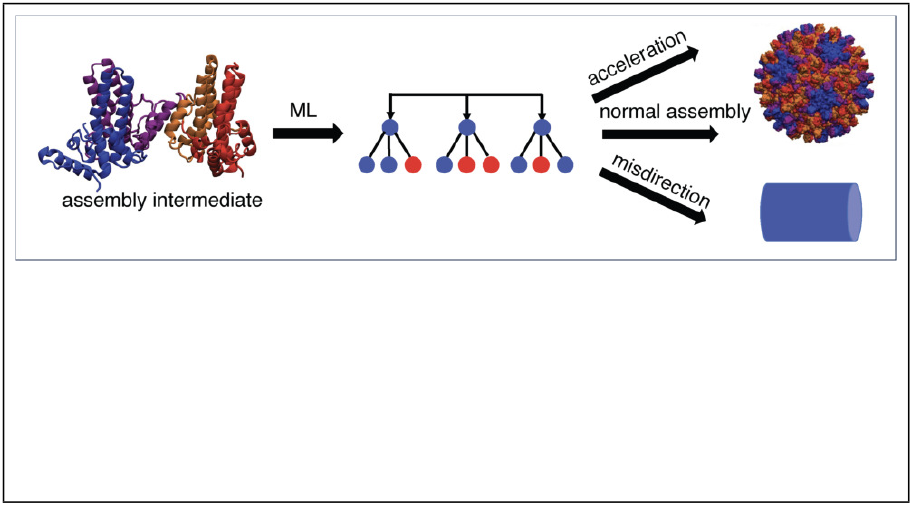

## Introduction

Hepatitis B virus (HBV) is one of the leading causes of liver failure and liver cirrhosis. Although vaccines against this virus are available, around 300 million people already suffer from chronic infections, making a cure, or at least active viral suppression, highly desirable.^1^ A novel promising approach is to target HBV nucleocapsid assembly, using compounds called capsid assembly modulators (CAMs). CAMs are typically small drug-like molecules that inhibit the viral life cycle by interfering with regular nucleocapsid assembly. Recent studies have shown that they can inhibit formation of covalently closed circular DNA (cccDNA) of HBV, which is responsible for persistent infections.^2–12^

The dominantly assembled HBV capsid has triangulation number T = 4 and is composed of 240 core protein (Cp) monomers (Figure 1a), although the assembly of T = 3 capsids with 180 copies of Cp is also possible.^13,14^ In the T = 4 capsid, Cp can adopt four quasi-equivalent conformations; A, B, C or D, depending on its position in the capsid (Figure 1a), with the DCBA tetramer (Figure 1b) forming the smallest building block. In the capsid, Cp can form either AB or CD dimers, which are structurally similar, with larger structural differences observed for the four quasi-equivalent dimer-dimer contacts.^13,14^

**Figure 1:**
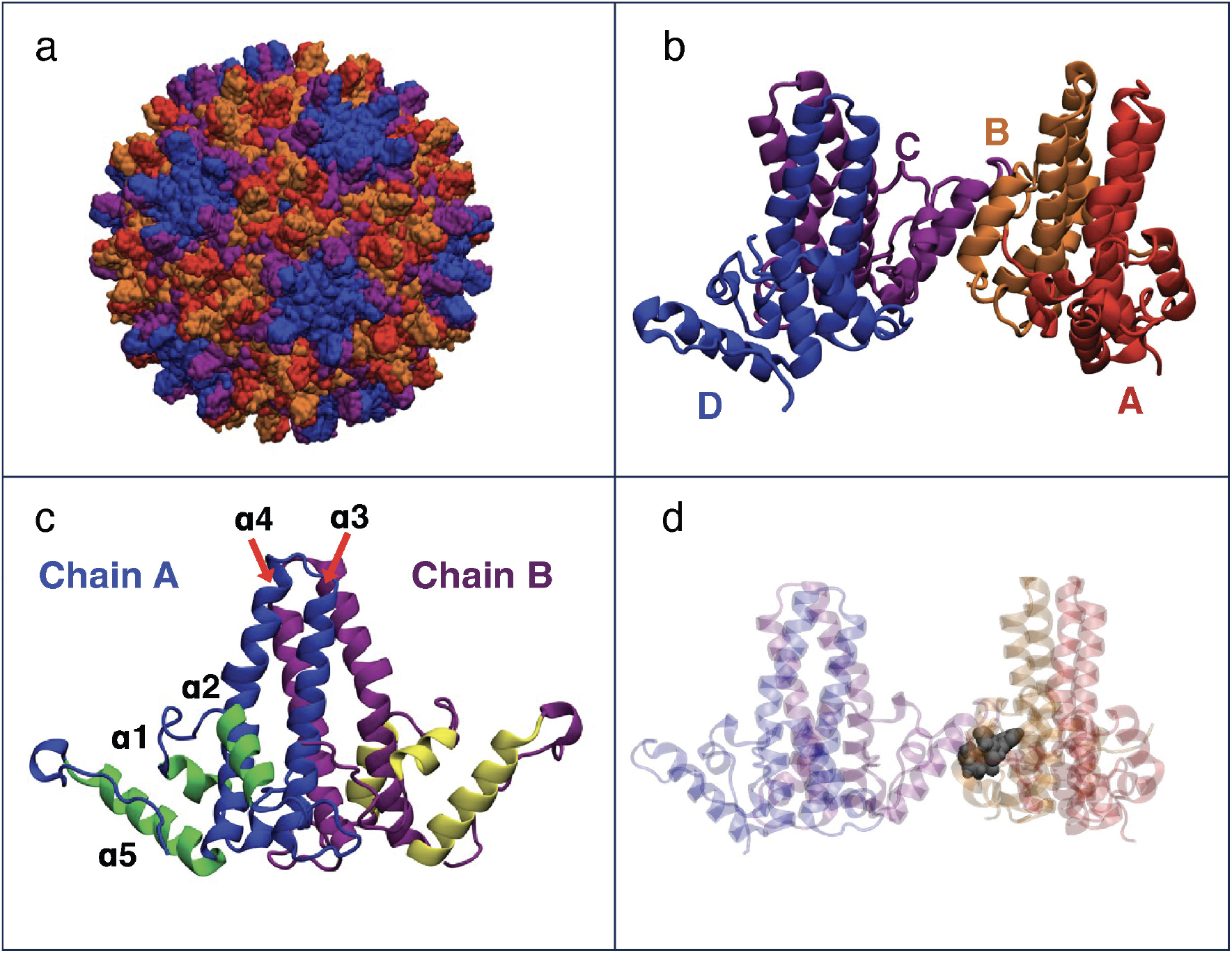
a) Structure of the complete T4 HBV capsid (coloring of the Cp149 proteins given in B). b) Cp149 tetramer, the building block of HBV capsids. The four distinct chains, A, B, C, D are shown in red, orange, purple, and blue, respectively. c) Structure of the Cp149 dimer, displaying helices α1-5. The spike region of the two monomers is shown in blue and purple for chain A and chain B, respectively, whereas the interface region is shown in green and yellow. d) Structure of Cp149 tetramer with bound CAM GLS4, shown in a gray space-filling representation, illustrating the binding site of all CAMs.

Pre-assembly, Cp exist as dimers, which can oligomerize into larger intermediates in a step-wise manner until a complete capsid is formed.^15,16^ The C-terminus of Cp is responsible for interacting with viral DNA, whereas the N-terminal domain forms the viral capsid.^17–19^ It has been shown that the N-terminal domain of Cp (Cp149) is sufficient to reproduce the capsid structure and capsid assembly mechanics. ^17,18^ Therefore, many assembly studies employ Cp149 in their experiments and simulations, including the MD simulations reported here.^5–7,12,20–25^ The Cp149 dimer consists of a spike region (helices α3 and α4), which protrudes outwards in the assembled capsid, and an interfacial region (helices α1, α2, and α5), which is responsible for inter-dimer interactions (Figure 1c). ^13,14^ Almost all known CAMs have been shown to bind at the interface between two dimers (Figure 1d).^2–7,9,10^ However, despite similar binding, a number of different assembly effects have been observed. Class I compounds cause the assembly of non-spherical structures, e.g., sheets or tubes,^2,3^ while in contrast, class II compounds cause the assembly of either regular or misshapen capsids.^4,5^ Some of the known chemical classes of CAMs are heteroaminopyrrolidines (HAPs) of class I^2,3^ and phenylpropenamides of class II^2,3^ (Figure S1).

Several experimental and computational studies have investigated the structural changes induced by different classes of CAMs. Hydrogen-deuterium exchange (HDX) experiments show that exchange in the Cp149 dimer increases upon binding of HAP18 and decreases in the assembled capsid, suggesting that HAPs destabilize hydrogen bonding in dimers and stabilize it in capsid.^26^ The most dramatic changes in HDX are observed for the top spike region and the α5 helices at the assembly interface. In addition, NMR experiments have shown changes in chemical shifts in the presence of both HAP compounds and CAMs based on sulfamoylbenzamide and glyoxamide derivatives for several Cp149 residues near the CAM binding site.^27^ Previous MD simulations have shown altered dimer-dimer orientation and C-terminus dynamics with bound HAP compounds in both capsids and early assembly intermediates.^24,28^ In our previous work, we studied dimers of dimers (tetramers) and trimers of dimers (hexamers), the first intermediates formed during assembly.^25^ Distinct inter-dimer orientations were observed in Cp149 tetramers and hexamers depending on the assembly effects of the bound CAMs. These differences could be represented by so-called base and spike angles (Figure 2b), which describe the opening/closure and the bending of the Cp149 tetramer, respectively.^25^ In addition we demonstrated the importance of correct CAM classification for rational drug design.^12^ Nevertheless, the specific details of how different assembly effects could be induced based on which compound type is bound to Cp have remained elusive.

**Figure 2:**
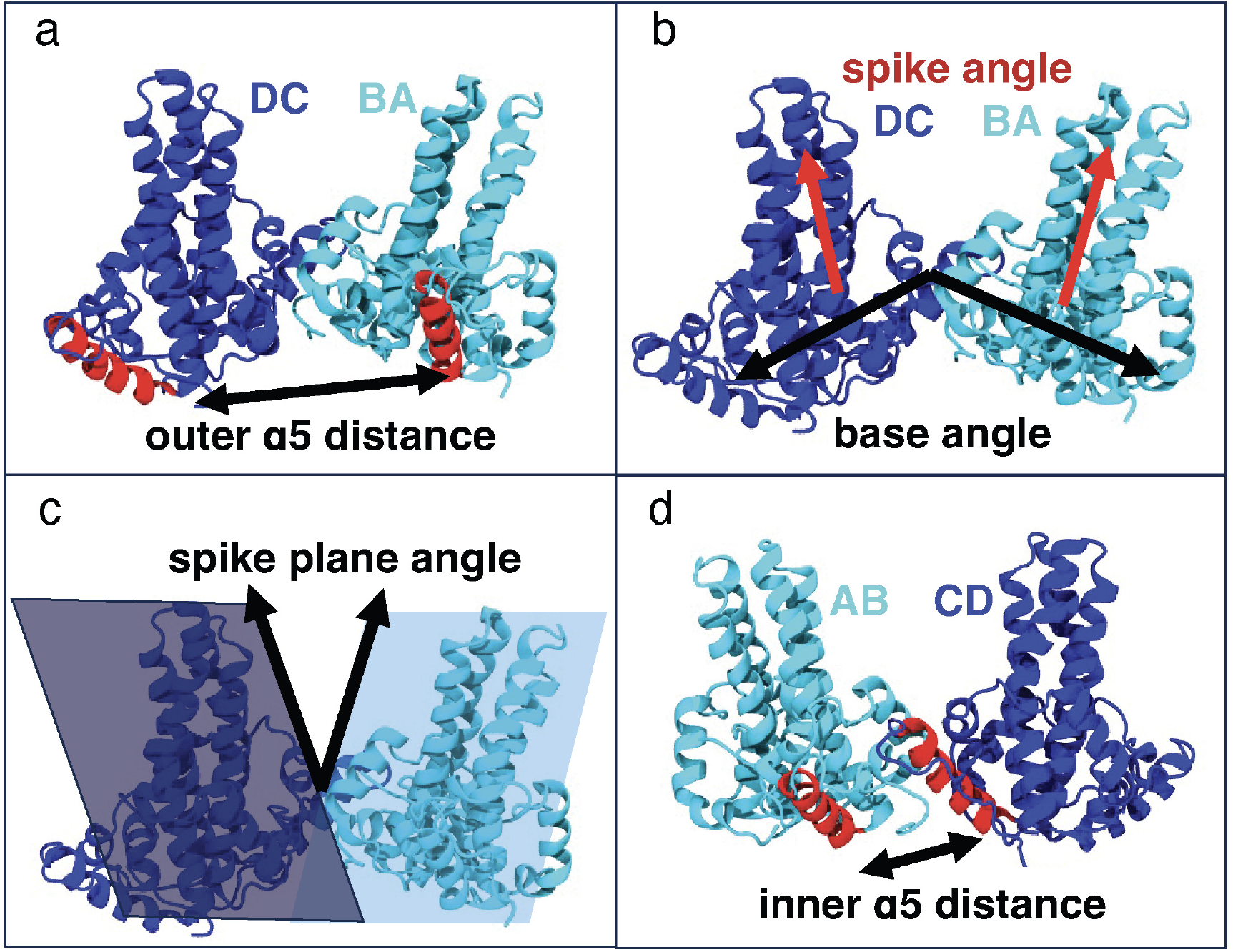
Illustration of the most important variables in our intuitive model. a) Outer α5 distance. b) Spike and base angles. c) Spike-plane angle. d) Inner α5 distance.

Previous work has shown that classification ML methods (CMLMs) can be particularly helpful for identifying differences between MD systems,^29–34^ and tools such as Scikit-learn^35^ and PENSA^32^ have been developed to help implement various ML approaches. However, to date, CMLMs or other ML approaches have not been applied to MD trajectories of HBV Cp. Notably, Fleetwood et al. applied several CMLMs to identify the most important features for conformational changes or ligand binding in specific systems, showing that the optimal method can depend on the studied problem.^29^ In addition, we previously applied ML with three different CMLMs to compare binding of SARS-CoV and SARS-CoV-2 S protein to its receptor ACE2, identifying several residues that contribute to distinct binding differences.^30^

In this work we combine MD simulations and ML to study HBV capsid tetramers with and without bound CAMs possessing different assembly effects. Specifically, HAP compounds (class I), AT130 (class II), and the V124W assembly-accelerating mutation are studied. Because the output of ML is sensitive to the input features used, we used different types based on tertiary structure or residue-residue distances. In addition, we tested four CMLMs, investigating their differences in accuracy and in selecting the most important features. The results for different input features and ML methods are compared in order to provide guidance for applications of CMLMs to other biomolecular systems.

## Methods

### System preparation

We used the following PDB structures as a starting point for each simulation: 3J2V^36^ for wild-type (WT) and V124W mutated systems, 4G93^37^ for AT130-bound system, and 5E0I^38^ for HAP1- and GLS4-bound systems. Table 1 also summarizes which structure was used for each system. 5E0I was crystallized with bound NVR-010-001-E2, which was modified into GLS4 or HAP1 with VMD’s Molefacture^39^ plugin for the corresponding systems, similar to our previous work^25^ (Figure S1). The missing C-terminal loops for each structure were added using ModLoop.^40,41^ The solvate VMD plugin was used to add explicit aqueous solvation to all systems, and the autoionize VMD plugin was used to add 0.15 M NaCl.^39,42^

**Table 1:**
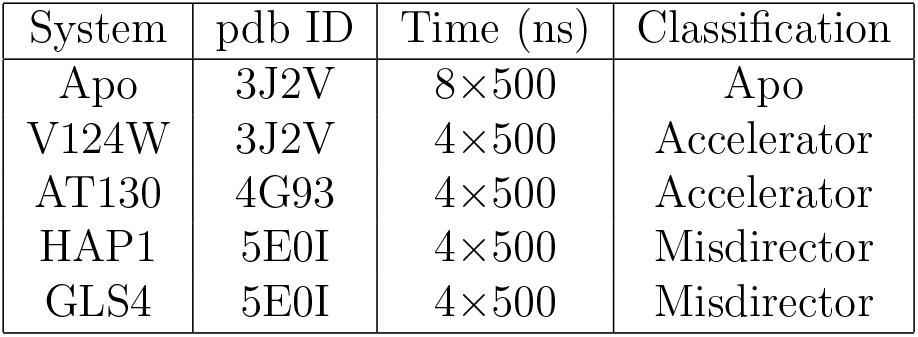
List of simulations used to build and validate our ML models.

| System | pdb ID | Time (ns) | Classification |
| --- | --- | --- | --- |
| Apo | 3J2V | 8×500 | Apo |
| V124W | 3J2V | 4×500 | Accelerator |
| AT130 | 4G93 | 4×500 | Accelerator |
| HAP1 | 5E0I | 4×500 | Misdirector |
| GLS4 | 5E0I | 4×500 | Misdirector |

### MD simulations

For all simulations, we used the CHARMM36m force field^43^ for the capsid protein and the TIP3P model for water.^44^ CGenFF^45^ parameters from the CGenFF webserver^46,47^ were obtained for the HAP1 and GLS4 molecules, whereas for AT130, we used previously optimized parameters.^48^

For the apo systems (WT and V124W), the energy of all atoms was minimized at once prior to MD simulations; however, for the systems with bound compounds (AT130, HAP1, and GLS4), a two-step minimization was performed instead. We first minimized the energy of water and ions only, followed by an energy minimization of all atoms. In our previous work, we observed that compound stability in the binding pocket can be increased with this two-step minimization.^49^ After minimization, we also performed a two-step equilibration for all systems. First, water and ions were equilibrated for 0.5 ns while the protein and the CAM (if present) were restrained. Next, the restraints were removed from the CAM and protein side chains, while the protein backbone was still restrained for 1 ns of equilibration. We used harmonic force constants of 2 kcal·mol^−1^·Å^−2^ for all restraints. In all MD simulations, we used rigid bonds for all hydrogen atoms, which allowed us to integrate the equations of motion with a 2-fs time step. A cutoff of 12 Å was used for the van der Waals interactions, with a smoothing function applied from 10 to 12 Å, ensuring a smooth decay to zero. For long-range electrostatic interactions we employed the particle-mesh Ewald method.^50^ The simulations were performed in the NPT ensemble, keeping the temperature and the pressure at the biologically relevant values of 310 K and 1 bar, respectively. The Langevin thermostat and the Langevin piston^51^ with a period of 200 fs and a decay of 100 fs were used to control the temperature and the pressure, respectively. The production runs were 500-ns long, and the first 50 ns were excluded from feature extraction in order to allow for system equilibration.

### Feature extraction

For the intuitive model, we used TCL scripts in VMD^39,42^ to obtain all of the features from the trajectories. For the angle and distance models, we switched to MDAnalysis^52^ in order to incorporate the large increase in the number of features. The distances and angles between parts of protein were calculated as follows. For distances, the geometrical center of the whole helix, or the lower or upper part of the helix, was measured for the backbone atom coordinates. The distances were then measured between helices or helix parts. Table S4 shows the residue definitions for each helix. For angles, each helix was split into a lower and an upper part, containing equal numbers of residues. The centers of the backbone atoms for each part were determined as in the distance case, allowing us to represent each helix as a vector. The angles between helices were obtained by calculating the angles between the helix vectors. For the distance model, the closest distance between any of the atoms in the residues was determined. In order to limit the number of features in this model, we only included the residue distances that were within 8 Å in at least one of the starting structures. We used a frame rate of 1/ns to extract features.

### Machine learning

All input features were normalized prior to their utilization in the ML models. We evaluated four distinct models: Logistic Regression (LR), Support Vector Machine (SVM), Random Forest (RF), and Multilayer Perceptron (MLP). The dataset comprised data points labeled as “apo”,”accelerator”, or “misdirector”, based on their corresponding simulation groups. To mitigate issues arising from multicollinearity, features exhibiting a correlation coefficient above 0.9 were identified using a correlation matrix and subsequently excluded. To ensure the robustness of this process, the order of features was randomized before each correlation assessment, thereby minimizing the impact of feature ordering on the exclusion criteria. The performance of these models was assessed through a five-fold cross-validation approach. To counteract the potential bias due to correlation of consecutive data points in simulation trajectories, each trajectory was segmented into five contiguous parts. Each segment was used sequentially as the testing set in the cross-validation process (Figure S2).

All four machine learning models were implemented using the scikit-learn library,^35^ outlined in their respective sections in the SI. The tuning of hyperparameters for each model was performed through a grid search approach with cross-validation, which systematically worked through multiple combinations of parameter values, allowing us to fine-tune the models to achieve a balance between computational cost and optimal accuracy on the validation data. This method ensured that the model not only fits our training data but also performs effectively on unseen data, thereby achieving good generalization. The feature importance was calculated individually for each model across 50 training iterations, and the mean importance of each feature was subsequently determined by dividing the cumulative importance by the frequency with which the feature was incorporated into the model. These values were then normalized to fall within the range of 0 to 1. After normalization, the importance scores were ranked independently for each model. This approach ensures that the relative significance of each feature is assessed consistently within the context of its respective model. See SI for additional details on each method.

Given the variability in feature importance distribution across models, we chose to average the rankings rather than the raw importance scores. This approach reduces the influence of outliers and provides a balanced view of feature significance across all models, culminating in an aggregated ranking that reflects the overall importance of each feature. This comprehensive ranking methodology ensures a more equitable representation of feature relevance, mitigating the undue influence of disproportionately weighted features.

For the angle-based model, features consisted of the angles between pairs of helices. The importance score of each individual helix was determined by dividing the accumulated importance by the number of occurrences of that particular angle in the features. Similarly, in the distance-based model, where the features were defined by the distance between pairs of residues, the importance score for each individual residue was calculated by dividing the accumulated importance by the frequency of inclusion of that specific distance in the features. This approach allows for the assessment of the relative contribution of each helix and residue to the model’s performance, providing insights into the most critical structural elements for conformational differences across classes.

## Results

We aimed to build an ML model with three classifications: apo, accelerator, and misdirector. Five Cp149 tetramer systems were simulated with MD: apo WT referred to as Apo, apo with the V124W mutation known to accelerate assembly,^23^ and tetramers with the bound accelerator AT130, or misdirector HAP1 or GLS4 (Table 1). For WT, eight trajectories were generated, while four trajectories were used for all other systems, ensuring equal simulation time for all three classifications. All trajectories were 500-ns long, and the first 50 ns were excluded from training and validation. Root-mean-square displacement (RMSD) of the protein backbone was measured for all trajectories, and appears to reach a plateau for the majority of the simulations after 500 ns (Figure S3). Four machine learning methods were used: LR, SVM, RF, and MLP (see Methods). In addition, four input-feature models were tested. The first model is called the intuitive model and consists of a limited set of features describing tetramer tertiary structure. These features were manually selected by examining the trajectories (Figure S4). The second model is based on angles between all of the helices in the tetramer. Finally, the third and fourth models use distances and inverse distances between pairs of residues in the tetramer, respectively. For all models, the accuracy and the most important features were evaluated. The feature ranking was also averaged between all ML methods to determine the overall importance.

### Intuitive model

For this model, we selected 20 features (Figure S4) including the spike and base angle descriptors from our previous work. Moderate accuracy (80-86%) was observed for the two analytical ML methods (SVM and LR), while higher accuracy (90-95%) was found for RF and MLP (Table S1). The distance between the two outer α5 helices (Figure 2a) was ranked highest overall, and it was also the top feature for all ML methods except LR, where it was ranked second (Table 2). More variance in ranking between the methods is observed for other features (Tables 2 and S2). Additional features that were ranked in the top five across all four methods were the spike and base angles, spike-plane angle (a modified definition of the original spike angle), and the inner α5 distance (Figure 2).

**Table 2:**
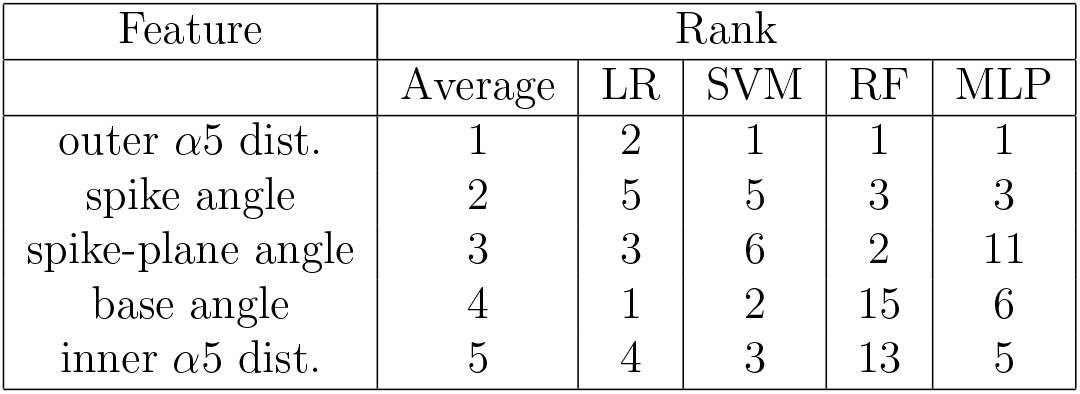
Ranking of top five important features across the four ML methods. Full ranking is shown in Table S2.

Whereas changes in the inter-dimer orientation described by base, spike, and spike-plane angles are noticeable by examining the trajectories, as in our previous work, ^25^ the changes in the α5 distances are less evident. However, distributions of the features in the trajectories show clear differences between simulated systems in all cases (Figure S5). In comparison to the Apo simulations, the inner α5 distance decreased and the outer distance increased with AT130 while the reverse was observed for the HAP compounds (Figure S5). The V124W mutation showed a similar inner distance to Apo and an outer distance smaller than in HAP-bound simulations. As in our previous work,^25^ increased spike and base angles were observed for AT130-bound and V124W systems, and decreased values are found for GLS4 and HAP1, with Apo system values ranging in between accelerator and misdirector values. The opposite changes in inner and outer α5 helix distances describe an inter-dimer movement similar to the base angle, with both movements corresponding to the opening or closing of the tetramer. Correlation analysis showed that the outer α5 distance is strongly correlated with the base angle, and the inner α5 distance is strongly anti-correlated with both spike and base angles (Figure S6).

While distributions of the top-ranked features (Figure S5) were different between accelerators, misdirectors, and Apo, significant overlaps between distributions were found, in particular between Apo, AT130-bound, and V124W systems, explaining the need for multiple features for accurate classification. In addition, differences were observed for some of the features for the AT130-bound and V124W systems, which are both considered accelerators, suggesting there may be differences in their mechanism of acceleration. The non-linear methods RF and MLP may perform better here because they can better account for the differences in features for AT130 and V124W. Due to the accuracy being lower than expected in comparison to our previous work where almost 100% accuracy was reached,^30^ we proceeded to test ML models that use more features.

### Angle-based model

In the angle-based model, all possible angles between the helices in the tetramer were calculated and used as input features. α3 and α4 helices were split in two because of the slight bending of these helices observed in many simulations. In total, each monomer in the Cp149 tetramer was divided into eight helices (Figure 3a). Each helix in the model is referred to by first its chain letter (A-D) followed by its helix number. α3 and α4 helices references also use a “t” or “b” at the end, depending on if the top or bottom part of the helix is indicated, respectively. Measuring all of the possible angles resulted in 496 features, which was reduced to ~350 features during training after removal of highly correlated variables (see Methods). High classification accuracy was observed for all ML methods, ranging from 96-98%, which was also a significant increase from the intuitive model (Table S1). In addition to the importance coefficients, which are obtained for specific helix-angle pairs, we also examined overall helix importance (see Methods). The latter approach captures the importance of each helix; however, it might overlook helices for which both very high and very low coefficients are present. Therefore, we compared the output of the most important helices and the most important helix pairs.

**Figure 3:**
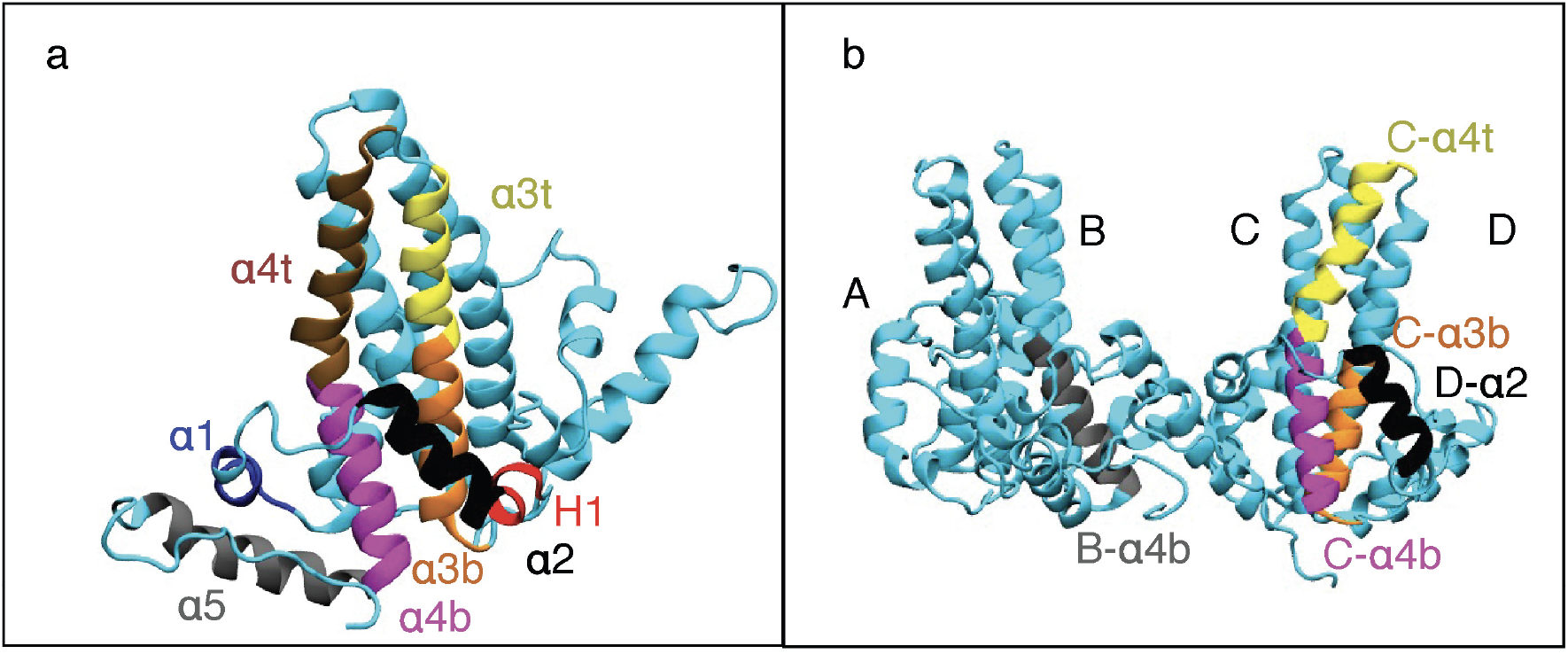
a) Division of Cp149 monomer into eight helices for our angle ML model. b) The top 5 most important helices according to our angle model, averaged over four ML methods. The importance order is B-α4b (grey), C-α4b (magenta), C-α4t (yellow), C-α3b (orange), and D-α2 (black). More data on importance is also shown in Table 3, and full data is provided in SI.

Examining the overall helix importance (Figure 3b and Table 3) and averaging the ranking across all four methods, we find that the spike region (α3 and α4 helices) is particularly important for classification, with four of the top five helices belonging to this region. Notably, B-α4b is the most important helix overall, and it is also the only helix from the spike region that interacts with the CAMs. Helices C-α4t and C-α4b are also highly ranked overall and across the four methods, in particularly for RF and MLP. Finally, visualization of helix importance (Figure 4) highlights the spike region for all methods.

**Table 3:**
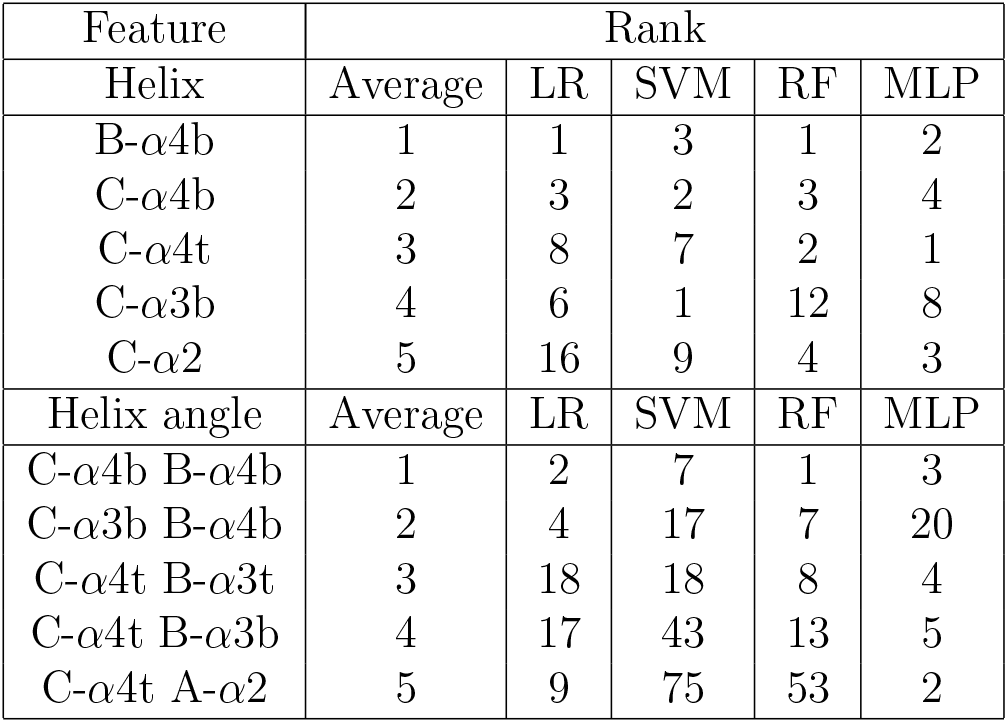
Ranking of top five overall important helices and helix angles.

| Feature | Rank |  |  |  |  |
| --- | --- | --- | --- | --- | --- |
| Helix | Average | LR | SVM | RF | MLP |
| B-α4b | 1 | 1 | 3 | 1 | 2 |
| C-α4b | 2 | 3 | 2 | 3 | 4 |
| C-α4t | 3 | 8 | 7 | 2 | 1 |
| C-α3b | 4 | 6 | 1 | 12 | 8 |
| C-α2 | 5 | 16 | 9 | 4 | 3 |
| Helix angle | Average | LR | SVM | RF | MLP |
| C-α4b B-α4b | 1 | 2 | 7 | 1 | 3 |
| C-α3b B-α4b | 2 | 4 | 17 | 7 | 20 |
| C-α4t B-α3t | 3 | 18 | 18 | 8 | 4 |
| C-α4t B-α3b | 4 | 17 | 43 | 13 | 5 |
| C-α4t A-α2 | 5 | 9 | 75 | 53 | 2 |

**Figure 4:**
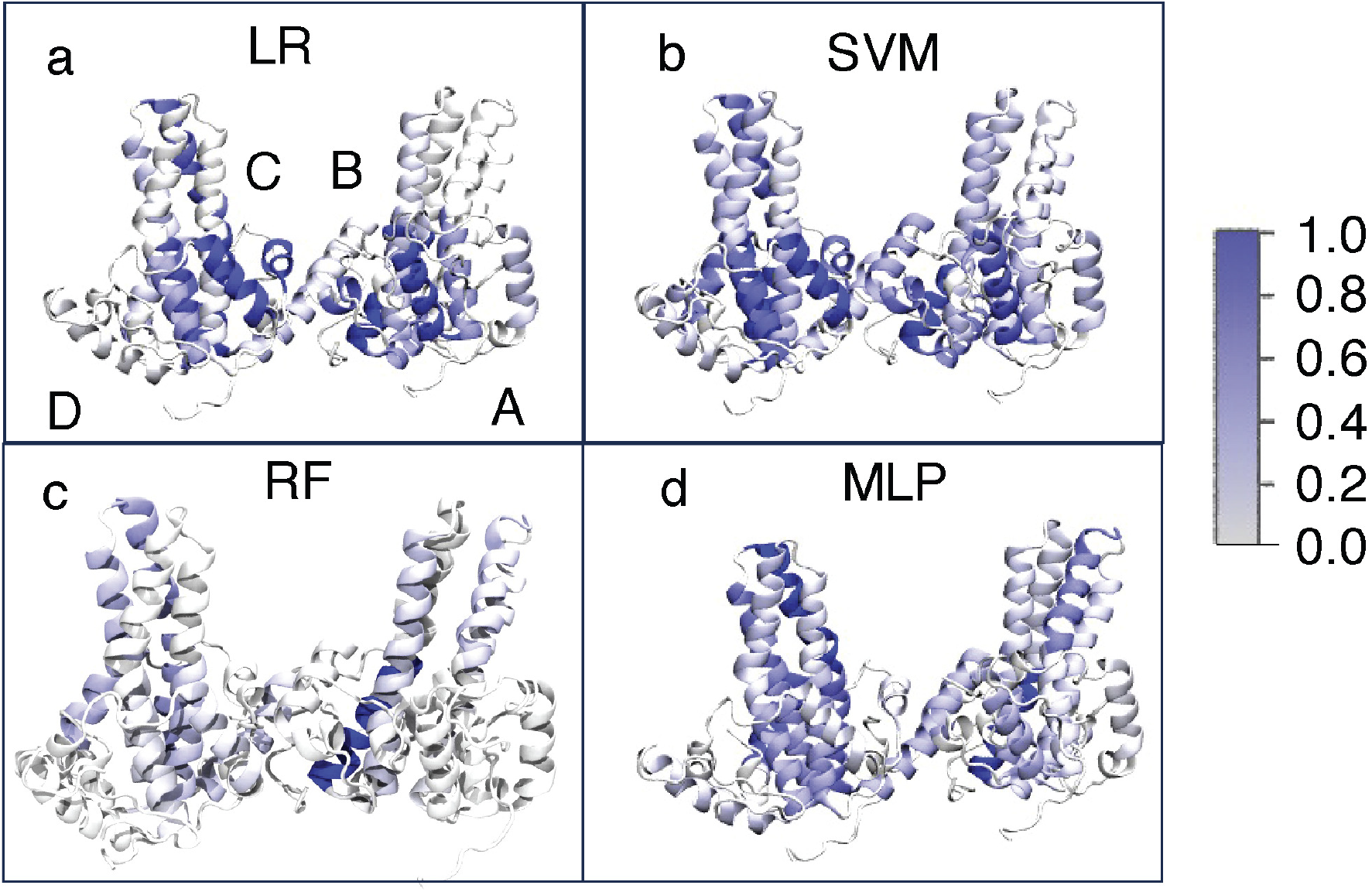
The importance profiles of our angle model for the four ML methods. The importance coefficients are scaled between 0 (no importance) and 1 (maximum importance). a) LR, b) SVM, c) RF, d) MLP.

Examining the most important helix pairs instead, we find that the most important angle is measured between the two top-ranked helices B-α4b and C-α4b (Table 3). Other important angles tend to be between at least one of the helices B-α4b, C-α4b, and C-α3b, and a spike region helix of the opposite dimer. The overall highly ranked helices are also frequently present in the highly ranked angles, reinforcing their importance. Structurally, the top angles have some similarity to the spike angle used in our previous work, which was measured between the bottom spike regions of each dimer. However, the differentiation between systems is greatly improved when the spike region is divided into multiple helices.

Examining the distributions for top five angles (Figure S7), we find most of them follow a similar trend to the spike angle, which measures the overall angle between spikes, with smaller angles for HAP1 and GLS4 (misdirectors), larger angles for AT130 and V124W (accelerators), and Apo angles in between the accelerators and misdirectors. Notably, for the C-α4b and B-α4b angle distribution in the V124W simulation appears more similar to the Apo system than the AT130-bound system, again suggesting differences between AT130 and V124W accelerator structures. Comparing the visualizations of helix importance for the four ML methods, we observe sparser importance profiles for RF, whereas more dispersed profiles are found for LR, SVM and MLP (Figure 4).

### Residue-distance model

The residue-distance model includes as features nearest-atom distances of all residues within 8 Å in at least one of the starting crystal structures. Because the C-termini are very flexible, all residues they could potentially interact with were included (see Methods). The number of initial input features was 13908, which was reduced to 8500 after exclusion of highly correlated features. A very high accuracy was obtained (>99% for all ML methods; Table S1), which could be attributed to a significant increase in the number of features compared to previous models. This sections focuses on the output of the model in which direct residue distances were used as input features, instead of inverse distances used in previous work.^29,30^ However, very similar results were obtained for the direct and inverse distances when comparing feature ranking (Tables 4 and S3). The overall order of highest-ranked residues is exactly the same for direct and inverse distances, although there are some minor differences in ranking for the individual methods. When examining top residues pairs instead, only minor differences in overall ranking and which residue pairs in the same region are included are observed. One notable difference was inclusion of the V124(C) and T109(B) residue pair in the inverse-distance model. Our initial analysis is focused on the output of the direct-distance model.

**Table 4:** Ranking of top ten overall important residues and residue pairs.

| Feature | Rank |  |  |  |  |
| --- | --- | --- | --- | --- | --- |
| Residue | Average | LR | SVM | RF | MLP |
| V124(D) | 1 | 1 | 1 | 2 | 6 |
| L140(B) | 2 | 5 | 12 | 3 | 2 |
| V76(A) | 3 | 2 | 2 | 1 | 20 |
| D2(A) | 4 | 3 | 4 | 9 | 31 |
| T142(B) | 5 | 16 | 9 | 28 | 1 |
| P79(A) | 6 | 22 | 3 | 7 | 34 |
| V93(C) | 6 | 7 | 17 | 10 | 32 |
| V93(C) | 8 | 24 | 15 | 13 | 17 |
| L140(D) | 9 | 20 | 36 | 11 | 8 |
| F18(D) | 10 | 49 | 10 | 16 | 4 |
| Residue pair | Average | LR | SVM | RF | MLP |
| V124(C) F103(B) | 1 | 2 | 2 | 1 | 2 |
| V124(C) L101(B) | 2 | 3 | 1 | 4 | 5 |
| V124(C) S106(B) | 3 | 7 | 6 | 2 | 1 |
| V124(C) C107(B) | 4 | 10 | 8 | 3 | 8 |
| G123(C) T109(B) | 5 | 11 | 28 | 5 | 4 |
| V124(C) F24(B) | 6 | 7 | 5 | 19 | 26 |
| V124(C) W102(B) | 7 | 4 | 4 | 17 | 33 |
| V124(C) I105(B) | 8 | 30 | 15 | 12 | 3 |
| V124(C) F110(B) | 9 | 15 | 25 | 16 | 13 |
| W125(C) F110(B) | 10 | 14 | 52 | 6 | 10 |

Examination of the top ten important residues across all ML methods (Figures 5a and S8 and Table 4) yields mostly hydrophobic residues spread between the spike region, C-termini, and interfacial region. None of these residues are part of C-α4b or B-α4b helices that were shown to be the most important ones by our angle-based model above. Also, only two of the top ten residues, L140(B) and T142(B), are in the vicinity of the CAM binding site. These two residues are expected to be affected by CAM binding as the backbone of L140(B) is known to interact with many CAMs.^25^ Inspection of the trajectories to identify the role of the remaining eight residues showed that the dissimilarities detected by ML likely originate in the differences between the starting structures used in this work (see SI for a detailed explanation). Examining the overall importance profiles between the four ML methods (Figure 6), we find very few residues with high importance coefficients for LR and RF and very disperse importance profiles for SVM and MLP, similar to the angle model. RF and LR only highlights a few residues close to the CAM binding region, helix α5 of chain D and top spike region of chain A. In contrast, SVM and MLP show significant importance in numerous regions of the Cp149 tetramer, although in comparison to the angle model, the CAM binding region and the outer interfacial region appear to be more important than the spike region.

**Figure 5:**
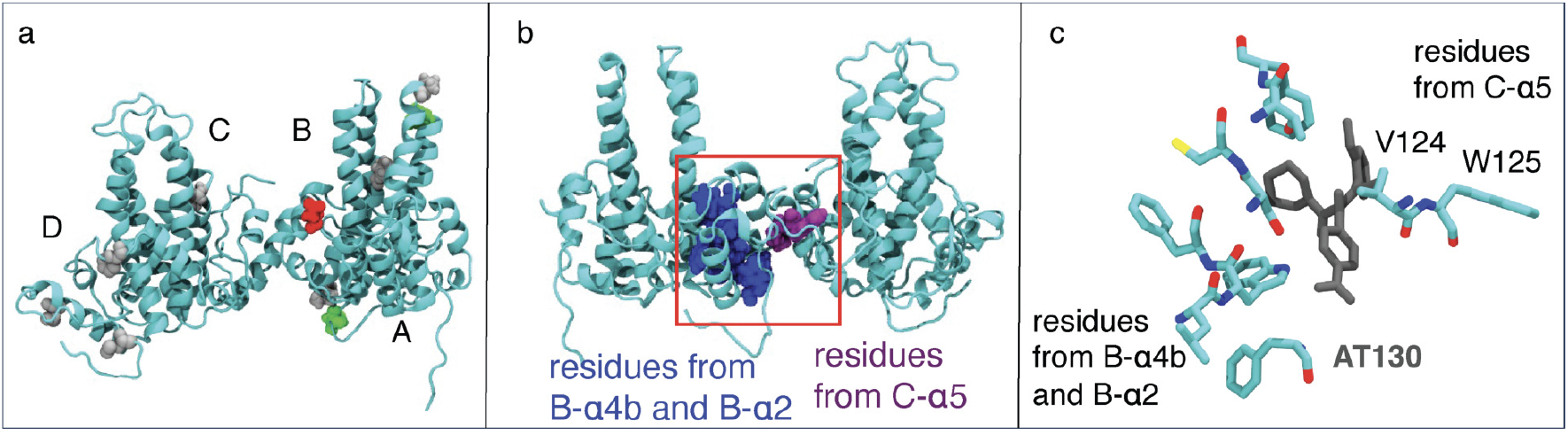
a) Top ten most important residues, averaged across four methods shown colored by residue type. Hydrophobic residues are colored grey, polar residues green, negatively charged residues red, and positively charged residues blue. A more detailed view of residues is shown in Figure S8. b) When analyzing the data from the top ten most important residue pairs, we mostly find distances between residues colored blue and purple. c) Detailed structure of the top ten residue pairs, illustrating the position of CAMs, in this case AT130, relative to the two residue groups.

**Figure 6:**
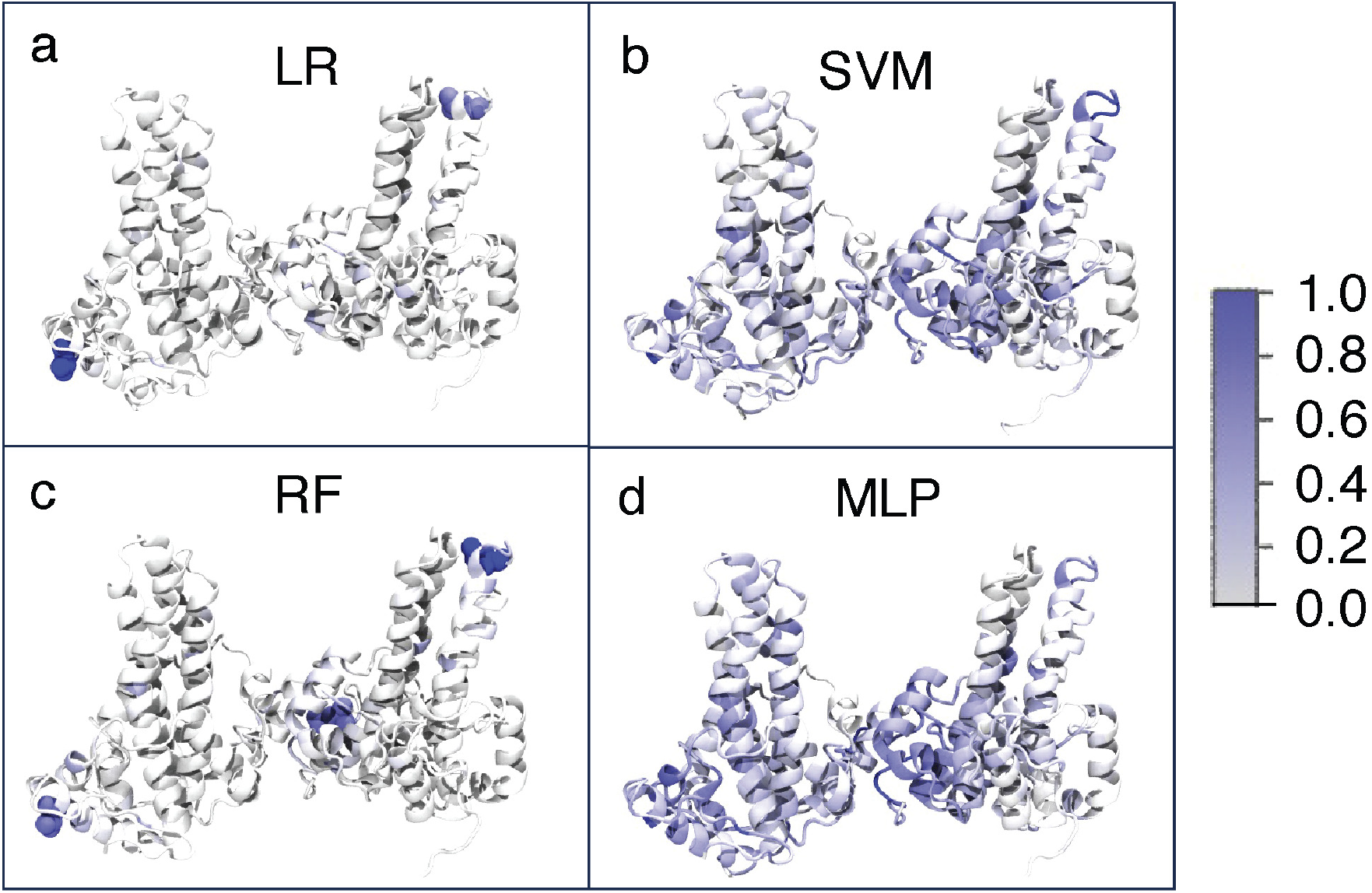
Importance profile of our residue-distance model for the four ML methods. The importance coefficients are scaled between 0 (no importance) and 1 (maximum importance). LR, b) SVM, c) RF, and d) MLP. In A and C, the residues with the normalized importance coefficient larger than 0.5 are also shown in space-filling representation.

When examining the top ten ranked residues pairs instead of top residues (Table 4), we mainly find various distances between α4b and α2 residues from chain B and α5 residues from chain C (Figure 5b). All of these residue are close to the CAM binding site, and many are in direct contact with the CAM. A closer look at the binding site (Figures 5b and S13) shows that the CAM binds right between these two groups of residues, which is expected to make these differences larger in the bound state. Examination of residue-distance histograms for the top-ten pairs (Figure S14) shows an increase in all distances for misdirectors HAP1 and GLS4 in comparison to both Apo and AT130 and a decrease for almost all distances involving residue W124 for the V124W system. The latter is expected due to the increased size and ability to form dispersion interactions for tryptophan in comparison to valine. The large differences in distances between HAPs and AT130 and similarities for many distances for AT130 and apo were surprising considering both HAPs and AT130 bind in the same pocket but can be explained by the different shapes of the ligands. A re-examination of the trajectories based on the ML output showed that for the HAP compounds, the ester moiety is wedged between T109 and V124 residues, separating them, whereas AT130 only has a modest effect on this distance in comparison to the apo state (Figures S13, S15, and S16). In addition, HAPs are bulkier molecules whereas AT130 is a longer molecule, which could explain larger distances between V124(C) and several residues in chain B for HAPs in comparison to AT130 (Figures S13 and S16).

## Discussion

MD simulation is a crucial tool for investigating differences between biological systems. However, such simulations produce a large amount of high-dimensional data, complicating the identifications of changes of interest. Furthermore, some important system dissimilarities may be overlooked because they are subtle or are not found where they are expected. Because ML approaches can analyze several thousand variables at once, they can be helpful in such cases. Notably, CMLM approaches can be used to classify new data after training on a dataset as well as identify which features are most important for the classification. It has been shown that the latter aspect of CMLMs can be utilized in MD simulations comparing different systems to identify the features that are most distinct between systems.^29–33^ Employing CMLMs to compare MD trajectories requires decisions on the method, input features for the model, and how to process the output. While previous studies have utilized inverse residue-residue distances within 15 Å and compared the output of different CMLMs after calculating the overall residue importance,^29,30^ we also compare the CMLM models based on different types of input features, and the importance of both overall and pair-wise features, if applicable. The different models are applied to HBV Cp149 tetramer bound with distinct CAMs bound, which can either accelerate the assembly or misdirect it into non-capsid structures. Our previous work identified subtle changes in the Cp149 tetramer structure based on the class of bound CAMs.^25^ In total, simulations with two classes of bound CAMs and one accelerating mutation are compared to the apo state. CMLM models were developed using four distinct methods (LR, SVM, RF and MLP), and three types of input features, a small user-selected dataset (intuitive model), a more comprehensive tertiary structure set composed of various helix angles (angle model), and a comprehensive set based on residue distances (residue-distance model).

Comparing the outputs for different types of input features, we found the intuitive model somewhat lacking in accuracy when classifying trajectory frames, with accuracy ranging from 83-93% (Table S1). The most important features were related to tetramer base and spike angles described previously.^25^ The angle model has significantly higher accuracy (up to 97%), likely due to its larger number of features, and highlighted several helices in the spike region as important, ranking the spike helix that also forms contacts with the CAMs (C-α4b) as the most important one (Figure 3 and Table 3). In contrast, the residue-distance model emphasized the importance of the CAM binding site and C-termini. The accuracy of this model was also further improved due to the addition of even more features, reaching up to 99%.

When comparing the two approaches of ranking for angle and distance models, we find significant agreement between highly-ranked helices and highly-ranked helix angles, especially among the top three features. However, some differences between the two lists are observed for lower-ranked helices (Table 3). The differences between pairwise or singular features are more pronounced for the residue-distance model. Ranking of the most important residues revealed a number of differences between the three starting structures that were used in our simulations (3J2V, 4G93, 5E0I). Because none of these observed changes are close to the CAM binding site, we theorize that these differences are structural artifacts due to lower resolution for some of the structures (Figures S9-12). In contrast, the pairwise feature ranking highlighted a number of residues at the CAM binding interface. Several residues from highly-ranked pairs were also shown to have increased NMR chemical shifts in presence of at least some CAMs: F24, W102, I105, S106, T109, and W125.^27^ Examining the distance distributions for the top ranked residue pairs (Figure S14) and the MD trajectories, we identify differences in binding of HAP compounds and AT130 (Figure S16). Notably, HAP compounds are bulkier, and their ester moiety is able to wedge between residues T109 and V124. It should be noted that residue pair V124(C) and T109(B) is not ranked in the top ten (rank 15); however, due to the prevalence of V124(C), T109(B), and F110(B) in the pair ranking, we focused our trajectory examination on these residues, discovering this difference. This pair was ranked higher (number 8) in the inverse-distance model, suggesting inverse distances might be more suitable for finding subtle changes in the trajectories. Previous work has also suggested that inverse distances are more suitable for detecting local changes.^29^

Comparing the output of the four ML methods, it is found that the rankings typically agree between the highest-ranking features (top 2-4), with more diversity between methods found for lower-ranked features. We hypothesize that any of the compared ML methods will pick up on particularly obvious features, whereas more subtle differences might be missed by some methods. Using more than one ML method should provide a more comprehensive picture of the differences. It is also found that SVM and MLP provide more disperse importance profiles, whereas more concentrated profiles are found for LR and RF. In our previous work comparing SARS-CoV and SARS-CoV-2 receptor binding, we found similar increases in dispersion and more sensitivity for the MLP method.^30^

The main novel finding from all of our models is the change in distances for several residues at the CAM binding site due to the bulkier HAPs, and the position of HAPs’ ester moiety between V124(B) and T109(C). Although T109 and V124 are not in direct contact in the crystal structure of the apo state (PDB code 3J2V), frequent contacts are observed in the MD simulations (Figure S15). Less frequent contact between these residues has been observed in previous simulations of whole HBV capsids,^24^ and some differences in residue contacts are expected when comparing capsids and smaller intermediates. The presence of AT130 only slightly increases the distances between these residues, while the presence of HAP1 or GLS4 causes a much greater increase (Figure S15). Mutations of T109 have been shown to confer HAP resistance and an increase in the number of normal capsids in the presence of HAPs,^24^ suggesting that this residue may play an importance role in assembly modulation. In addition, increased NMR chemical shifts for T109 have been observed in the presence of several CAMs.^27^ More experimental studies on the effects of T109 mutations on the assembly in the presence of different CAM classes are needed to clarify the effects of this residue.

Our work shows that models based on tertiary-structure features are more suitable for detecting large-scale allosteric changes in comparison to a residue-distance based model, which was focused on detecting minor differences between residues. This is expected because a distance cutoff is typically required in a residue-distance model in order to maintain a manageable number of features, which limits the model’s ability to detect large-scale changes. Our work also illustrates the power of CMLMs in detecting structural differences in MD trajectories that can easily be overlooked by a human observer as well as the need for visual inspection of trajectories to clarify the reasons for the obtained output from the models. The described approaches could be used to classify novel HBV CAMs, and compare their modes of actions as well as study other biological systems of interest.

## Abbreviations

HBV: hepatitis B virus
Cp: capsid protein
CAM: capsid assembly modulator
MD: molecular dynamics
cccDNA: covalently closed circular DNA
HAP: heteroaryldihydro pyrimidine
PPA: phenylpropenamide
SBA: sulfamoyl benzamide
NMR: nuclear magnetic resonance
HDX: hydrogen-deuterium exchange
ML: machine learning
WT: wild type
RMSD: root-mean-square displacement
CMLM: classification machine learning method
SVM: support vector machine
LR: logistic regression
RF: random forest
MLP: multilayer perceptron

## Acknowledgement

This work was supported by the National Institutes of Health (R01-AI148740). We also used the Hive cluster in this work, which is supported by the National Science Foundation (MRI- 1828187) and is managed by the Partnership for an Advanced Computing Environment at Georgia Tech.

## Supporting Information Available

Additional analysis including 16 figures and four tables.

## Notes

### Competing Interest Statement

The authors have declared no competing interest.

