## Supporting Information for "Machine learning of molecular dynamics simulations provides insights into modulation of viral capsid assembly"

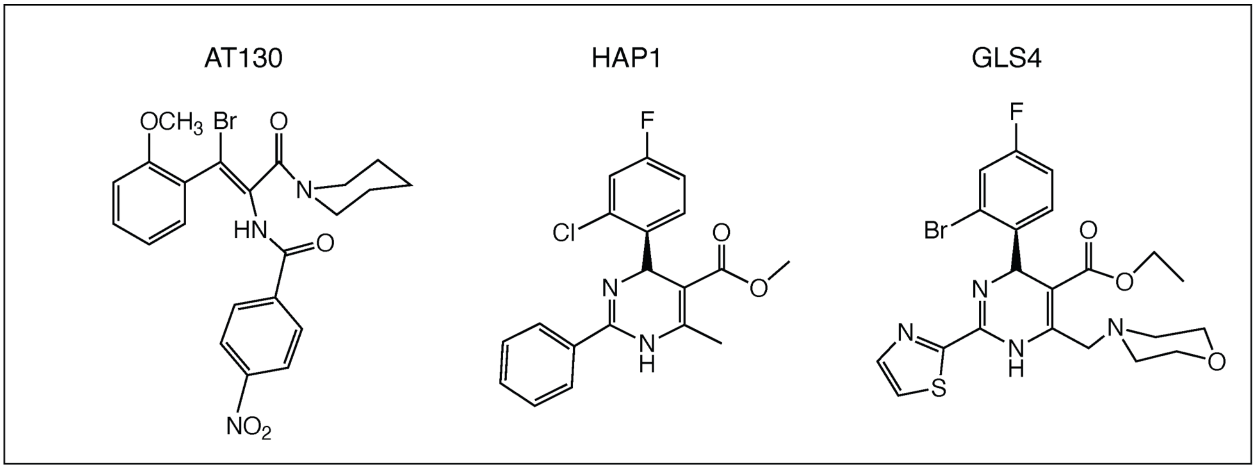


Figure S1: Structures of CAMs studied here.

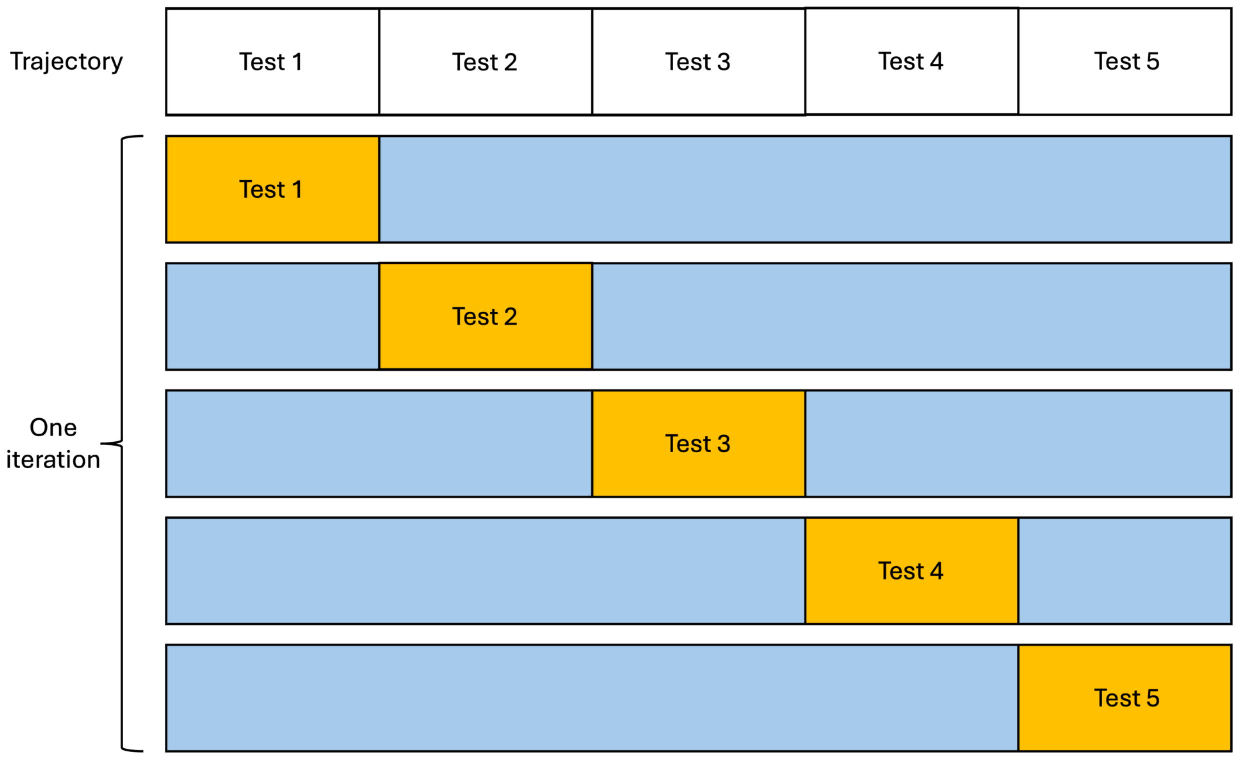

Figure S2: The setup of cross validation in our machine learning models.

Table S1: Classification accuracy, as fraction of correctly classified frames, for all employed feature sets and methods.

| Feature set | Method | | | |
|---|---|---|---|---|
| | LR | SVM | RF | MLP |
| Intuitive | 0.834 | 0.836 | 0.932 | 0.945 |
| Angle | 0.974 | 0.973 | 0.958 | 0.982 |
| Distance | 0.993 | 0.997 | 0.997 | 0.999 |
| Inverse distance | 0.995 | 0.998 | 0.995 | 0.9997 |

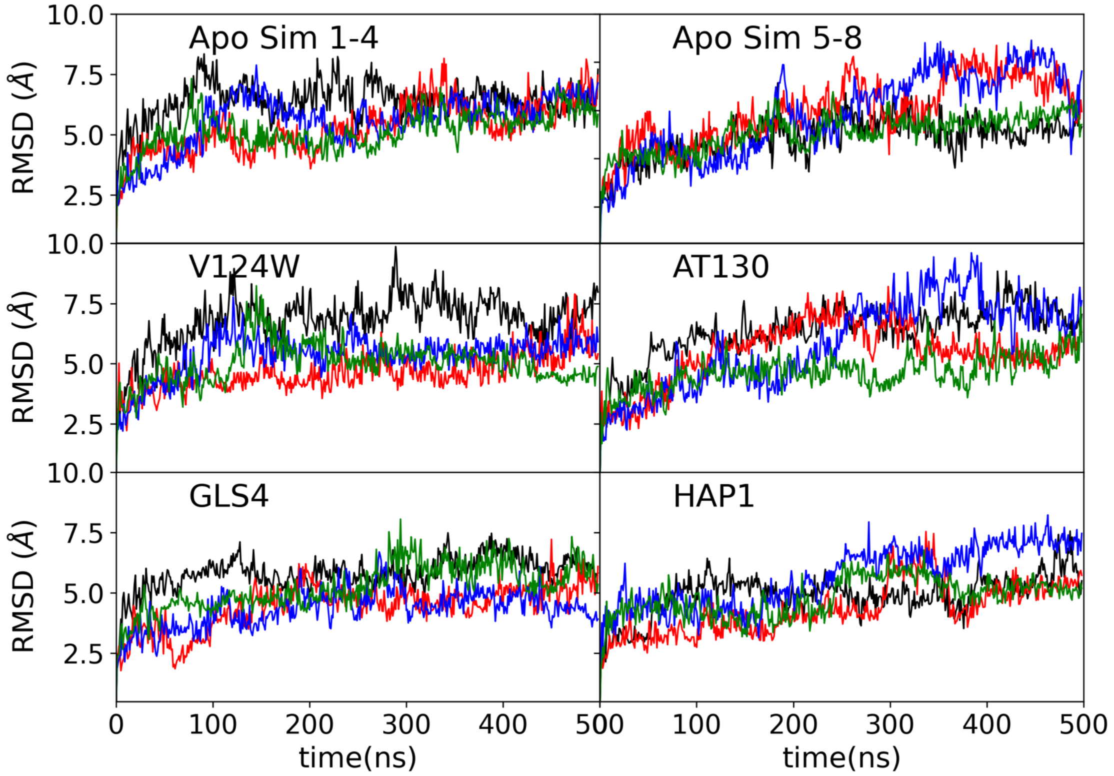


Figure S3: RMSD graphs for all of the systems, the different colors correspond to independent runs.

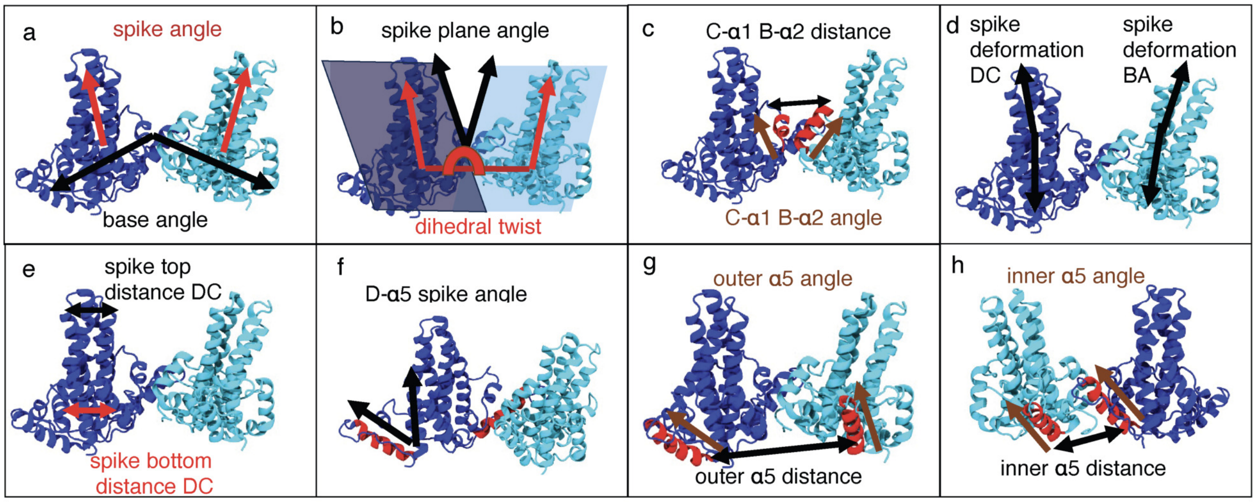


Figure S4: Features selected for the intuitive ML model. a) Base and spike angles used in our earlier work. b) Spike plane angle and dihedral twist variables. Spike plane angle is similar to the spike angle, however, here the plane going through each dimer is determined based on the position of the $\alpha5$ helices, and the angle is measured between the two planes. For the dihedral twist variable we construct and measure the dihedral angle between the top part of the spike region for the two dimers in the tetramer. c) Angle and distance between C-$\alpha1$ and B-$\alpha2$ helices, the distance was measured between the bottom part of the helices. Both helices are shown in red. d) Spike deformation angle, which measures the angle between the bottom and the top part of the spike. This angle is measured for AB and CD dimers. e) distance between the two monomers in the dimer at the top and bottom of the spike region. These distances are measured for AB and CD dimers. f) Angle between D-$\alpha5$ (red) and the spike region. This angle is measured for each monomer. g) Distance and angle between the outer $\alpha5$ helices (red). The distance between the bottom part of the helices is measured. h) Distance and angle between the inner $\alpha5$ helices (red). The distance between the bottom part of the helices is measured.

Table S2: The ranked importance of all features of the intuitive set.

| Feature | Rank | | | | |
|---|---|---|---|---|---|
| | Average | LR | SVM | RF | MLP |
| outer $\alpha5$ dist. | 1 | 2 | 1 | 1 | 1 |
| spike angle | 2 | 5 | 5 | 3 | 3 |
| spike-plane angle | 3 | 3 | 6 | 2 | 11 |
| base angle | 4 | 1 | 2 | 15 | 6 |
| inner $\alpha5$ dist. | 5 | 4 | 3 | 13 | 5 |
| C $\alpha2$ B $\alpha1$ dist | 6 | 9 | 9 | 9 | 2 |
| CD bottom spike dist. | 7 | 8 | 4 | 4 | 15 |
| A spike $\alpha5$ angle | 7 | 10 | 11 | 6 | 4 |
| dihedral twist | 9.0 | 11 | 8 | 7 | 9 |
| C $\alpha2$ B $\alpha1$ angle | 10 | 12 | 12 | 5 | 8 |
| AB botton spike dist. | 11 | 7 | 7 | 8 | 18 |
| CD top spike dist. | 12 | 15 | 15 | 11 | 10 |
| C spike $\alpha5$ angle | 13 | 13 | 17 | 10 | 12 |
| outer $\alpha5$ angle | 14 | 6 | 10 | 19 | 20 |
| inner $\alpha5$ angle | 14 | 14 | 13 | 14 | 14 |
| AB top spike dist. | 16 | 16 | 18 | 17 | 7 |
| B spike $\alpha5$ angle | 17 | 17 | 16 | 12 | 16 |
| AB deform. | 18 | 18 | 14 | 20 | 13 |
| CD deform. | 19 | 20 | 19 | 18 | 17 |
| B spike $\alpha5$ angle | 19 | 19 | 20 | 16 | 19 |

### Differences in starting crystal structures found from analysis of top ranked residues in the residue-distance model

The first difference that we found was that in all of the starting structures and Cp149 monomers except chain D of PDB 4G93, R133 and W125 are parallel stacked in the fold between $\alpha5$ and C-terminus (Figures S9c,d). However, vertical stacking is instead observed for chain D of the 4G93 structure used as a starting state for AT130 simulations (Figure S9b). The horizontal stacking is maintained in MD simulations, whereas vertical stacking is not (Figures S9e-g). The lack of stacking allows other residues to move in the $\alpha5$ helix fold, altering the distances between several residues in that region, including L140 and V124 (Figure S9e). Furthermore, in the mutated V124W system, W124 can form contacts with R133 or W125, increasing the importance of this residue and potentially others in the vicinity (Figure S9e). The structural difference described above is a likely cause for the high ranking of residues V124(D) and L140(D). The lack of stacking between R133 and W125 also propagates to nearby residues, affecting contacts of R127 and V126 with F18, which explains high ranking of F18(D) (Figure S10)

Next, we examine the residues of chain V76(A) and P79(A), finding that V76 is typically folded into the spike region, with the exception of chain A in the structure of 3J2V (apo), where it points toward the solvent instead (Figure S11). This initial structural difference remains throughout the MD simulations and alters the distances between V76 and other nearby residues, including P79. For residue 93, we find that it is a different amino acid for some of the structures due to slightly different HBV strains used in the experiments. This residue is M in the 3J2V structure of HBV subtype adw , and V in 4G93 and 5E0I structures of HBV subtype adyw.

Finally, examination of the N-terminus of chain A showed a different orientation of M1 relative to I3. In 3J2V chain A, the side chain of M1(A) points away from I3(A), and in all other cases it points toward I3(A) (Figure S12) This structural change is expected to affect several distances in the N-terminal region, with the ML methods finding D2(A) as the most impacted residue.

### Additional details on ML approaches

##### SVM model

SVM is designed to find the hyperplane that optimally separates the data into different classes by maximizing the margin, thus establishing a robust decision boundary. In our research, the SVM model, implemented using the scikit-learn library, classifies data points from MD trajectories into three categories: "apo", "accelerator", or "misdirector". These classifications are based on their normalized features.

Given that the standard SVM model inherently handles binary classification, we adopted the "One-vs-One" (OVO) strategy to manage multi-class classification challenges. This approach involves training an SVM classifier for each pair of classes, resulting in a total of $\frac{3(3-1)}{2} = 3$ individual classifiers for the three classes involved. When classifying a new sample, each of these classifiers casts a vote for one of the classes, and the class with the majority of votes is assigned to the sample.

To ensure simplicity and minimize the risk of overfitting, a linear kernel was utilized across all SVM models. The regularization parameter C, critical for controlling the trade-off between achieving low training errors and maintaining a sufficiently large margin, was fine-tuned using a grid search method. The grid included values of C such as 0.1, 1, 10, and 100. After comprehensive evaluation through cross-validation, the optimal value of C selected for the final models was found to be 1, which provided the best balance between complexity and generalization capability across our data sets.

Feature importance was assessed through the coefficients derived from these linear SVM models. Each coefficient in the SVM model represents the contribution of a feature to the decision boundary, allowing us to quantify the impact of each feature on the classification results.

##### RF model

The RF algorithm, an ensemble learning method, constructs multiple decision trees during training and predicts output based on the mode of the classifications from these trees. We implemented this algorithm using RandomForestClassifier in scikit-learn library to categorize data into three distinct classes based on features extracted from our dataset.

In fine-tuning the model, we adjusted several key hyperparameters to optimize both performance and computational efficiency. The number of trees in the forest was set at 500, balancing accuracy with processing time, and each tree was limited to a maximum depth of 50 to prevent overfitting. Furthermore, to control model complexity, we varied the number of features considered for the best split: 5 for intuitive model, 20 for angle model, and 100 for distance models.

Feature selection during training was critically based on the reduction of Gini impurity, which measures the decrease in uncertainty or disorder at each node of the trees. The final importance of each feature was determined by the cumulative reduction in Gini impurity contributed by the feature across all trees. This method highlights the most impactful features as well as enhances the overall accuracy and interpretability of the model by focusing on the most informative attributes.

##### LR model

LR is a fundamental tool for classification tasks, utilizing a logistic function to estimate the probabilities that each instance in the dataset belongs to a particular class. This function effectively maps any real number to a value between 0 and 1, ensuring that output values are interpreted as probabilities. In our research, we employed scikit-learn's LogisticRegression implementation to categorize data into three distinct classes: "apo", "accelerator", or "misdirector".

For our model, we applied L1 regularization, also known as lasso, which is instrumental in feature selection by shrinking less important feature coefficients to zero, thereby enhancing model interpretability and avoiding overfitting. The regularization strength was set with a value of 1, balancing the bias-variance tradeoff by providing a moderate level of regularization.

Feature importance in logistic regression was extracted by examining the coefficients of the model. The magnitude and sign of each coefficient indicate the strength and direction of the influence that a particular feature has on the probability of belonging to a specific class. Larger absolute values correspond to features that have a more significant impact on the model's decisions, allowing us to identify which variables are most influential in classifying molecular dynamics trajectories.

##### MLP model

MLP is an advanced neural network method that employs multiple layers of neurons to capture complex patterns in data. We implemented this model using MLPClassifier in scikit-learn to classify data into three distinct categories based on features derived from our dataset.

In tuning our MLP model, we carefully adjusted several key parameters to optimize both performance and computational efficiency. The network was configured with a single hidden layer with varied number of neurons: 5 for the intuitive model, 20 for the angle model, and 100 for the distance models, chosen based on the number of features for each model. The activation function used was ReLU (rectified linear unit), which is known for helping in accelerating the convergence of stochastic gradient descent and preventing the vanishing gradient problem.

For the optimization algorithm, we selected 'adam', renowned for its robustness and efficiency, particularly with large datasets. The learning rate was set to 'constant', ensuring steady progression in weight adjustments across iterations, and the model was allowed a maximum of 200 iterations to converge to a solution. This controlled approach to learning and iteration helps balance the need to train the model thoroughly against the computational expense of training.

To extract and understand the contribution of individual features to the model's predictions, we utilized Layer-Wise Relevance Propagation (LRP), which works by decomposing the MLP's output decision back to the input feature level, illuminating how each feature influences the overall prediction.

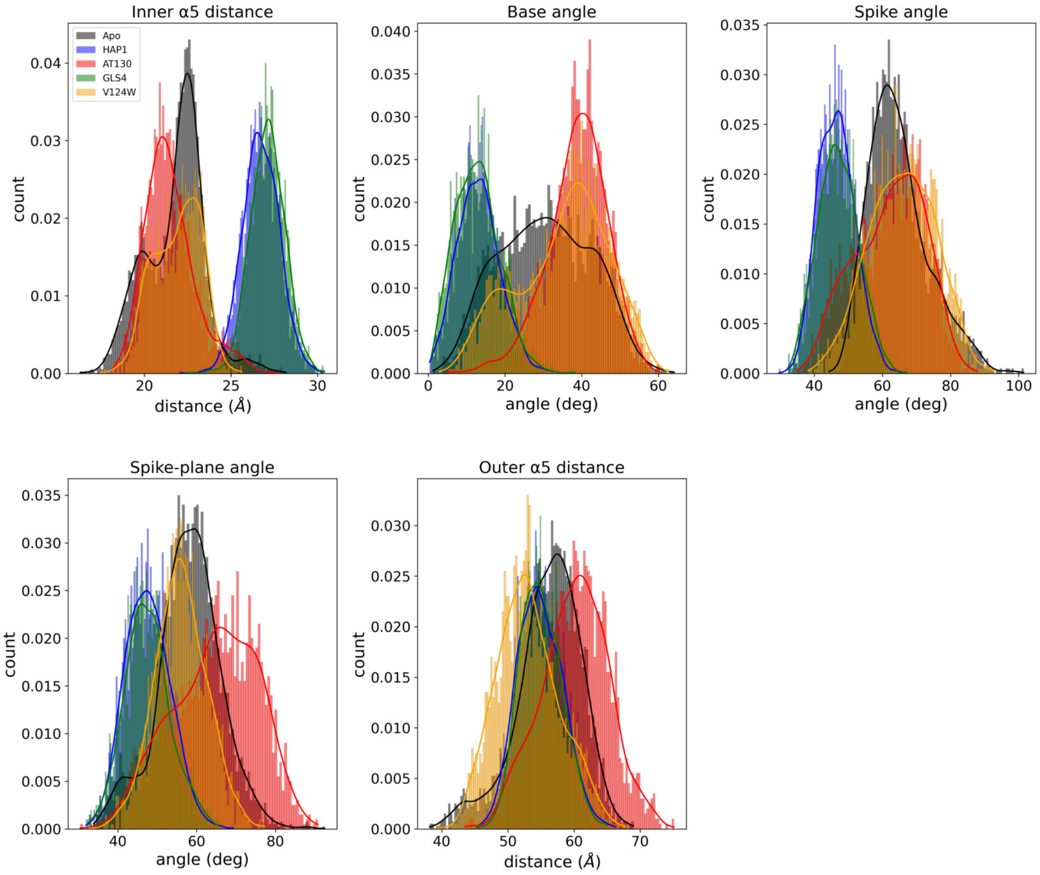


Figure S5: Distribution histograms of the five most important features for the intuitive model.

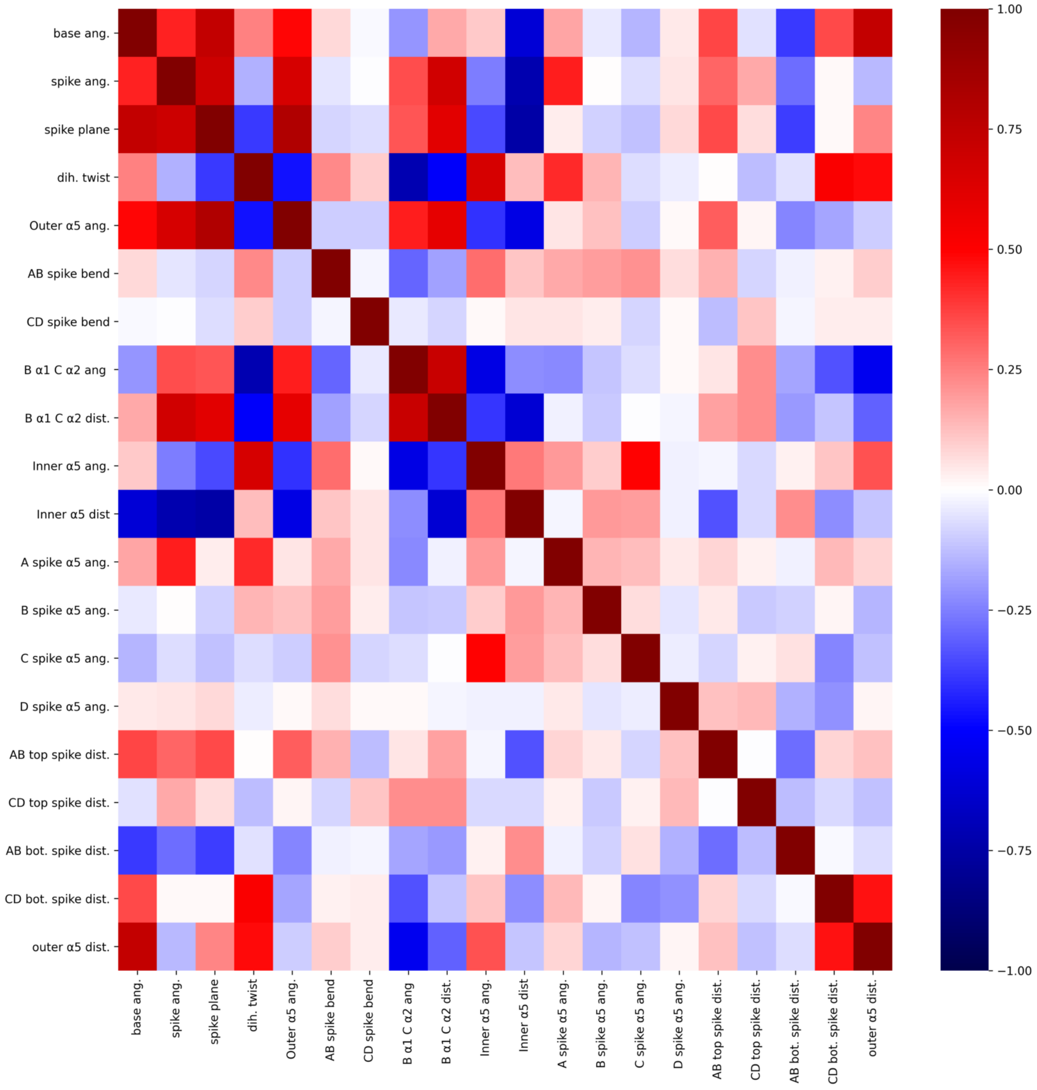


Figure S6: Correlation between the variables in the intuitive model.

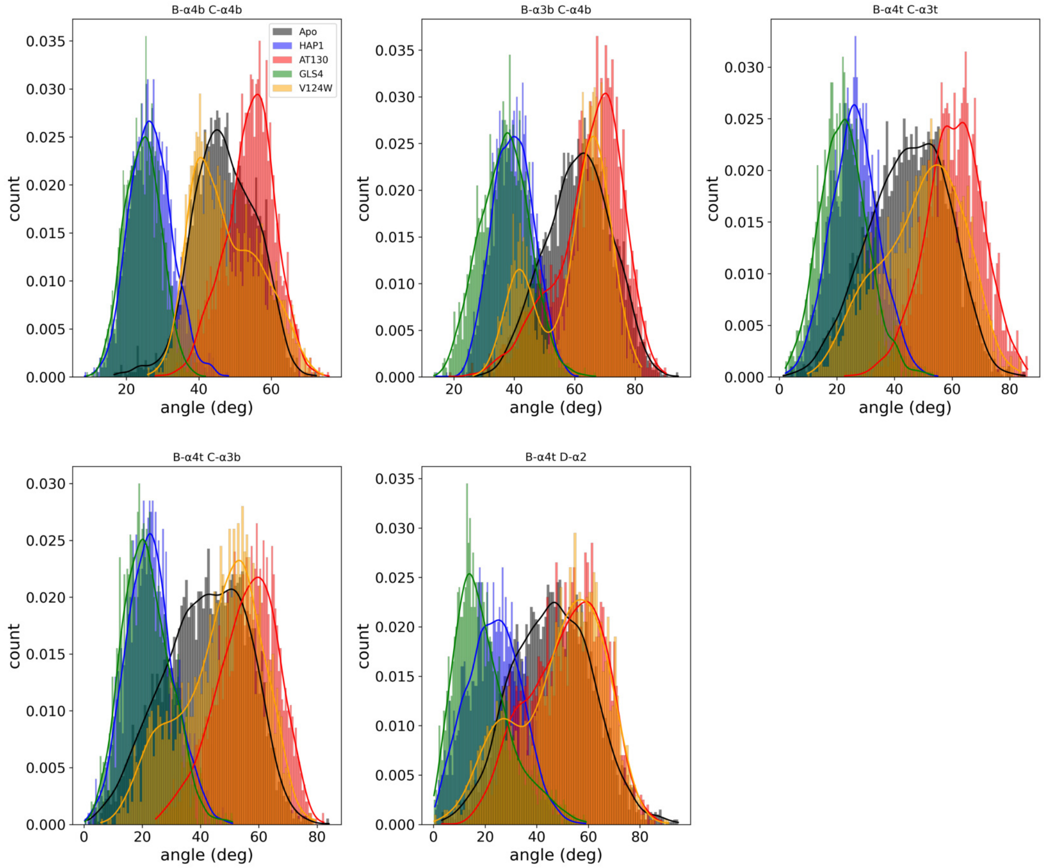


Figure S7: Distribution histograms of the five most important features for the angle model.

Table S3: The ranking of top residues and top residue pairs for the reverse distance model.

| Feature | Rank | | | | |
|---|---|---|---|---|---|
| Residue | Average | LR | SVM | RF | MLP |
| V124(D) | 1 | 6 | 1 | 2 | 1 |
| L140(B) | 2 | 2 | 12 | 3 | 5 |
| V76(A) | 3 | 20 | 2.0 | 1 | 2 |
| D2(A) | 4 | 31 | 4 | 9 | 3 |
| T142(B) | 5 | 1 | 9 | 28 | 16 |
| P79(A) | 7 | 34 | 3 | 7 | 22 |
| V93(B) | 7 | 32 | 17 | 10 | 7 |
| V93(C) | 8 | 17 | 15 | 13 | 24 |
| L140(D) | 9 | 8 | 36 | 11 | 20 |
| F18(D) | 10 | 4. | 10 | 16 | 49 |
| Residue pair | Average | LR | SVM | RF | MLP |
| V124(C) S106(B) | 1 | 4.0 | 3.0 | 2.0 | 1.0 |
| V124(C) L101(B) | 2 | 2.0 | 2 | 4 | 3 |
| V124(C) F103(B) | 3 | 1 | 1 | 1 | 13 |
| V124(C) C107(B) | 4 | 9 | 7 | 3 | 2 |
| V124(C) W102(B) | 5 | 3 | 4 | 15 | 43 |
| V124(C) A34(B) | 6 | 6 | 8 | 51 | 6 |
| V124(C) T109(B) | 7 | 12 | 18 | 17 | 29 |
| V123(C) T109(B) | 8 | 15 | 24 | 5 | 46 |
| V124(C) I105(B) | 9 | 62 | 11 | 12 | 9.0 |
| V124(C) F24(B) | 10 | 7 | 6 | 24 | 59 |

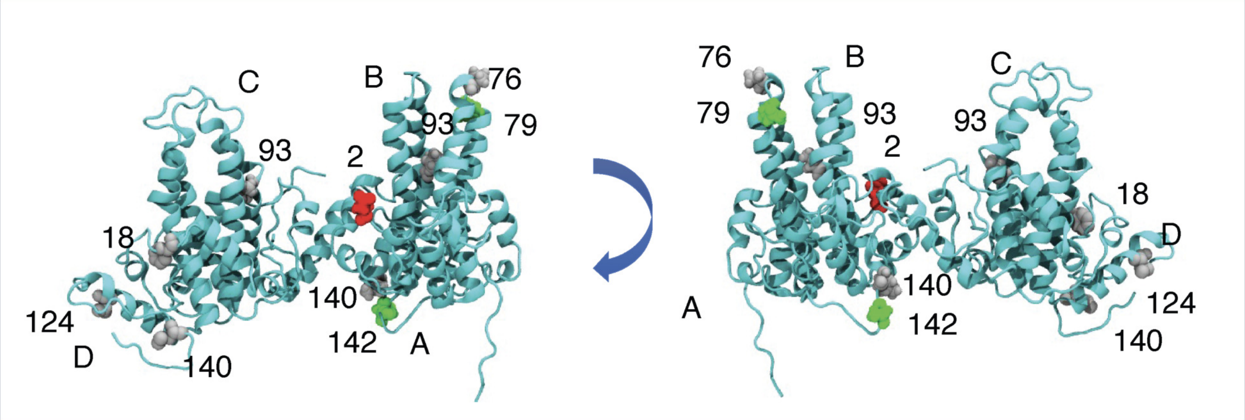


Figure S8: The top ten most important residues, averaged across four methods are shown colored by residue type. Hydrophobic residues are colored grey, polar residues are colored green, negatively charged residues are colored red and positively charged residues are colored blue. Both sides of the tetramer are shown for clarity, and residue numbers are added.

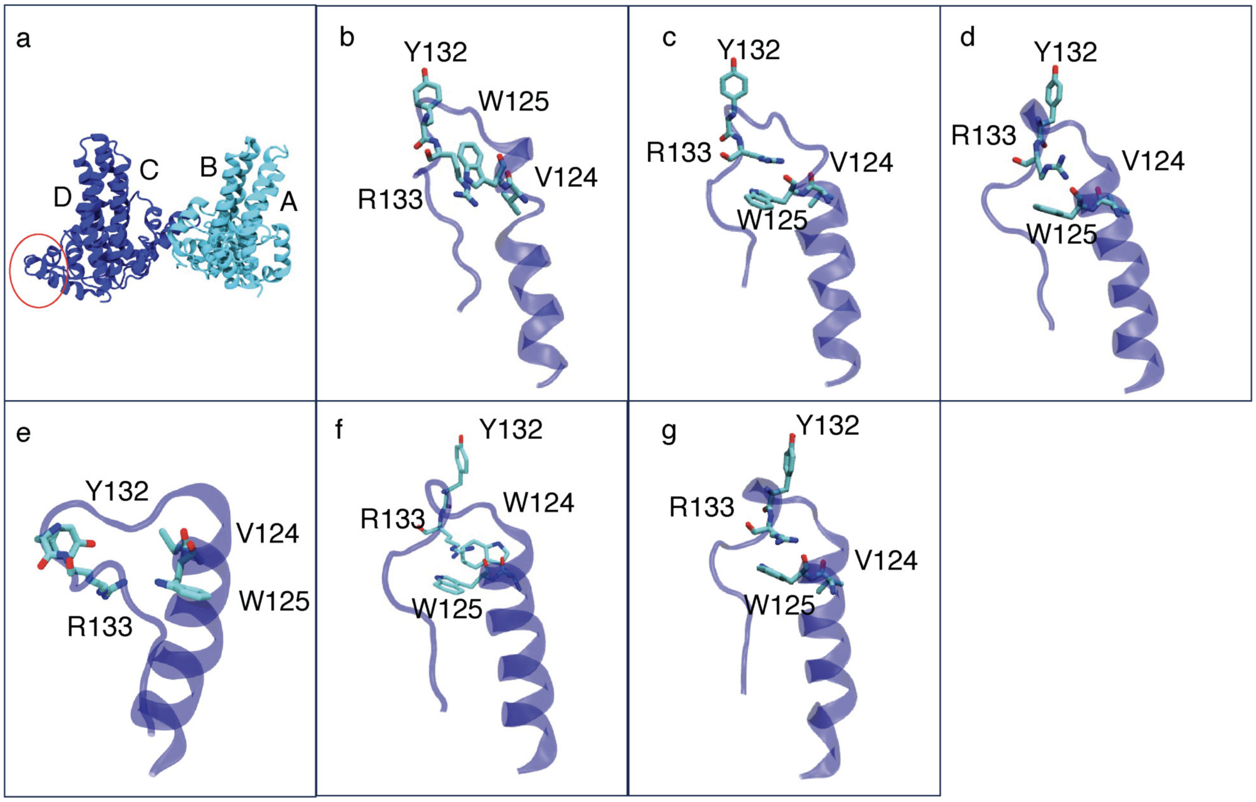


Figure S9: Comparison of the $\alpha 5$ C-terminal region between the used crystal structures and MD simulation. a) Figure of the full tetramer with the region enlarged in other subfigures marked by a red circle. b) Structure of D monomer in 4G93 (AT130-bound), notably R133 and W125 are stacked in a vertical orientation. c) Structure of A monomer in 4G93, notably R133 and W125 are stacked in a horizontal orientation. This orientation is also observed in monomers B and C of 4G93, and all monomers of 3J2V and 5EI0 structures. d) Structure of D monomer in 3J2V showing the parallel orientation of R133 and W125. e) A snapshot from AT130-bound simulations for, showing separation of R133 and W125 in chain D, which in turns alters the position of residues Y132(D) and W125(D). f) A snapshot from W124V simulations, showing retained stacking of residues R133(D) and W125(D), the position of Y132(W) is similar to the starting orientation, however, the mutated W124(D) is shown to form contacts with R133(D) and W125(D). g) A snapshot from Apo simulations, showing that the structure of residues of interest, R133(D), V124(D), W125(D) and Y132(D) does not change significantly from the starting crystal structure.

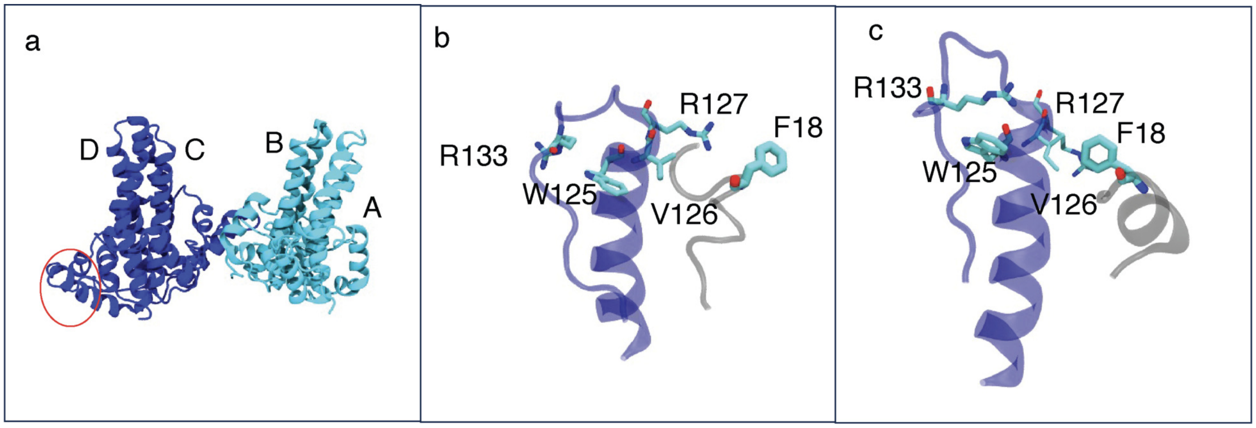


Figure S10: Comparison of the $\alpha5$ C-terminal region and the residues close to F18 for chain D in AT130-bound and Apo simulations. a) Figure of the full tetramer with the region enlarged in other subfigures marked with a red circle. b) A snapshot from AT130 simulations that shows how the separation of R133(D) and W125(D) also alters positions of R127(D) and V126(D) relative to F18(D). c) A snapshot of from Apo simulations showing that close contacts between R133(D) and W125(D) are retained, which in retains close contacts between F18(D) and R127(D) and V126(D).

Table S4: Residue numbers in Cp149 corresponding to each helix defined in our ML model.

| Helix | residue numbers |
|---|---|
| $\alpha1$ | 12-18 |
| $\alpha2$ | 27-39 |
| H1 | 40-44 |
| $\alpha$3b | 49-63 |
| $\alpha$3t | 64-75 |
| $\alpha$4t | 78-95 |
| $\alpha$4b | 96-110 |
| $\alpha5$ | 111-127 |

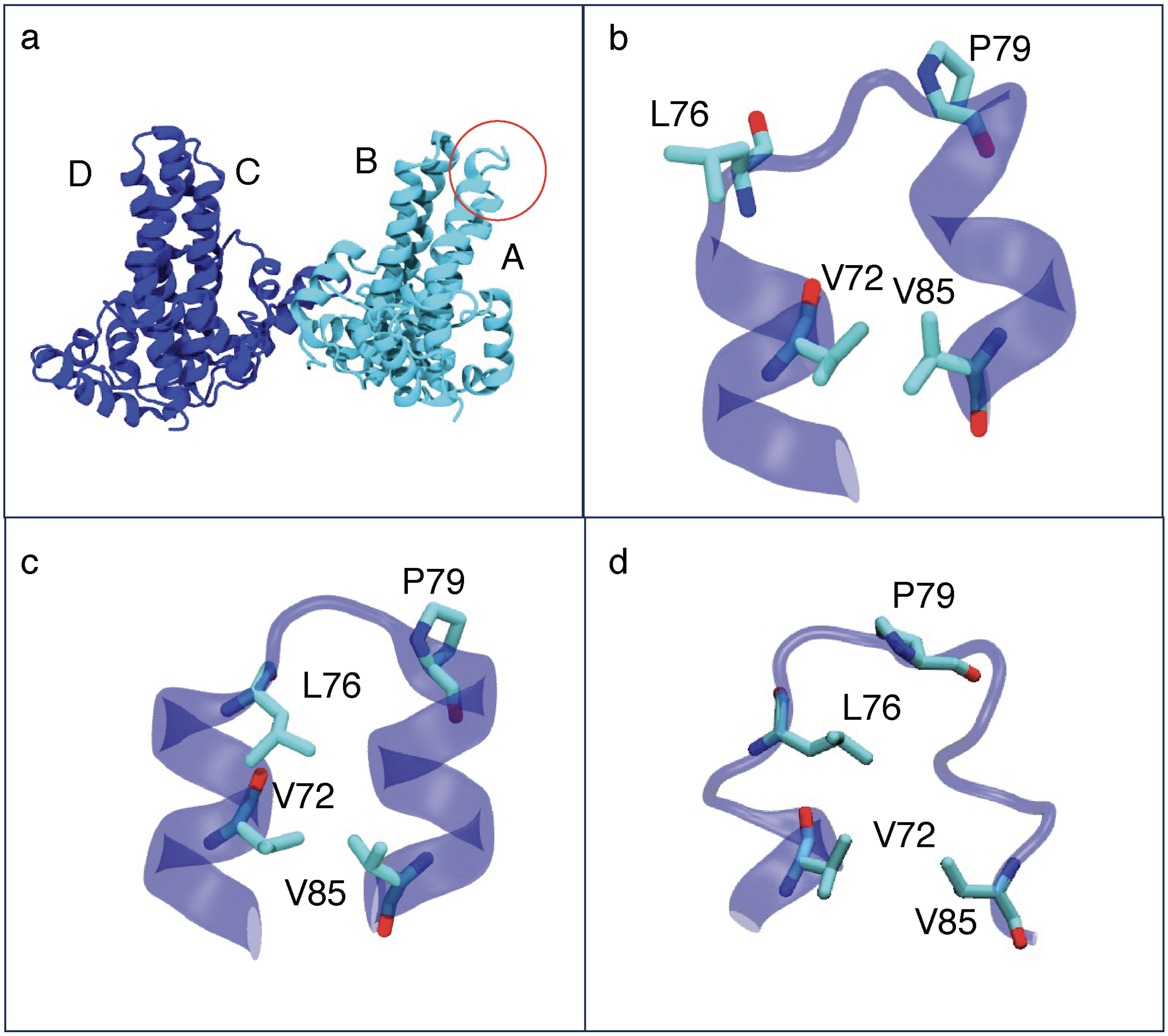


Figure S11: Comparison the of top spike region of the A chain between Apo and AT130 simulations. a) Structure of the full tetramer with the red circle marking the region enlarged in other subfigures. b) Structure of the top spike region for the A monomer in 3J2V (Apo). In particular the different orientation of L76(A) away from V72(A) and V85(A) is shown. c) Structure of the top spike region for the D monomer in 3J2V (Apo). The different orientation of L76(D), where it forms contacts with V72(D) and V85(D) is shown. This orientation is also found for monomer B and C. d) Structure of the top spike region for the D chain in 4G93 (AT130-bound). Although minor differences in P79 orientation are seen, the L76 side chain points inward, as in c.

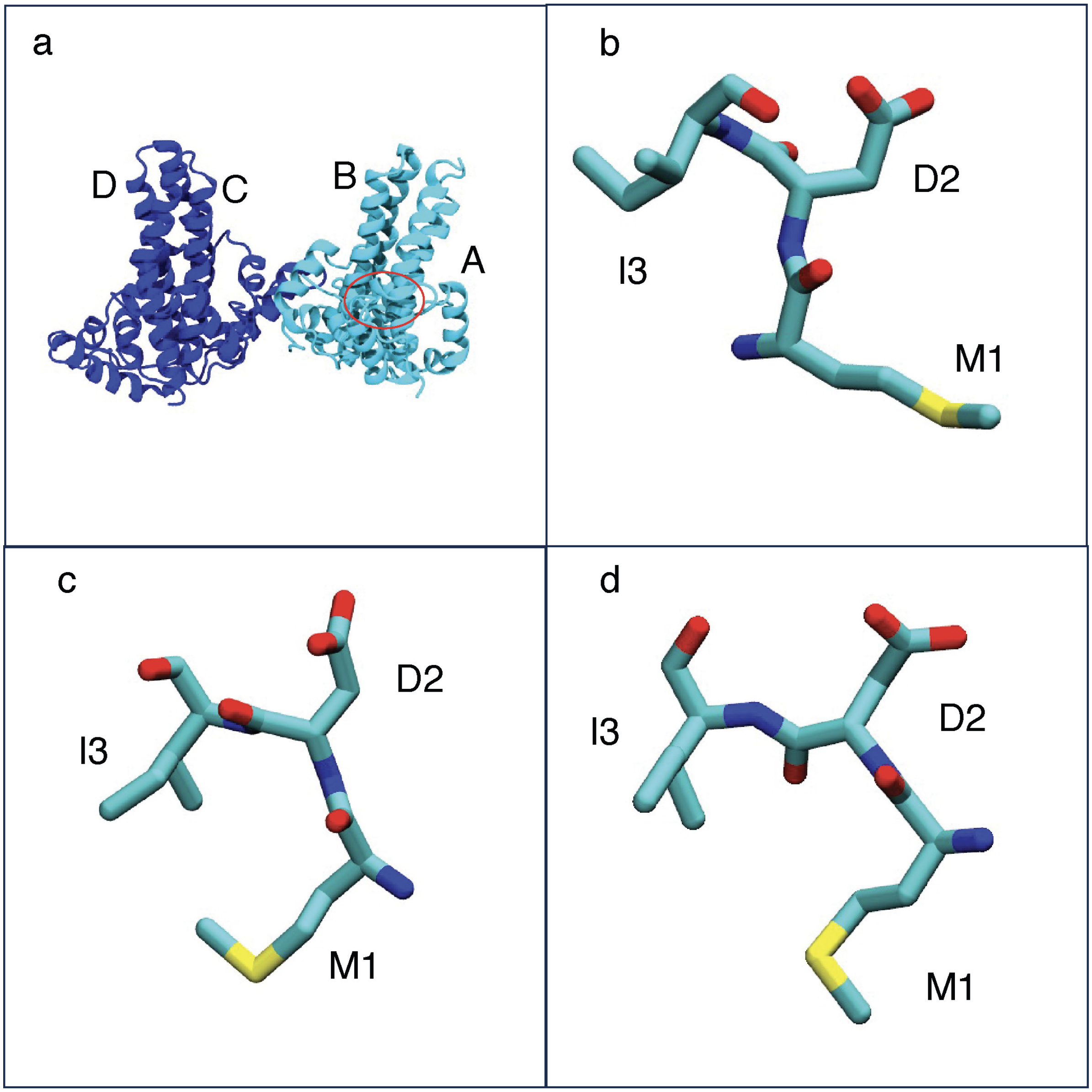


Figure S12: Comparison of N-terminal structure. a) Figure of the full tetramer with the red circle marking location of N-terminal of monomer A b) The conformation of the first three amino acids for monomer A in 3J2V (Apo) structure. Notably M1 side chain is shown to point away of I3. c) The conformation of the first three amino acids for monomer D in 3J2V (Apo) structure. In this case M1 side chain is shown to point towards I3. d) the conformation of the first three amino acids for monomer A in 4G93 (AT130-bound) structure. Although minor differences from 3J2V non-A monomers are seen, M1 side chain is still closely positioned to I3.

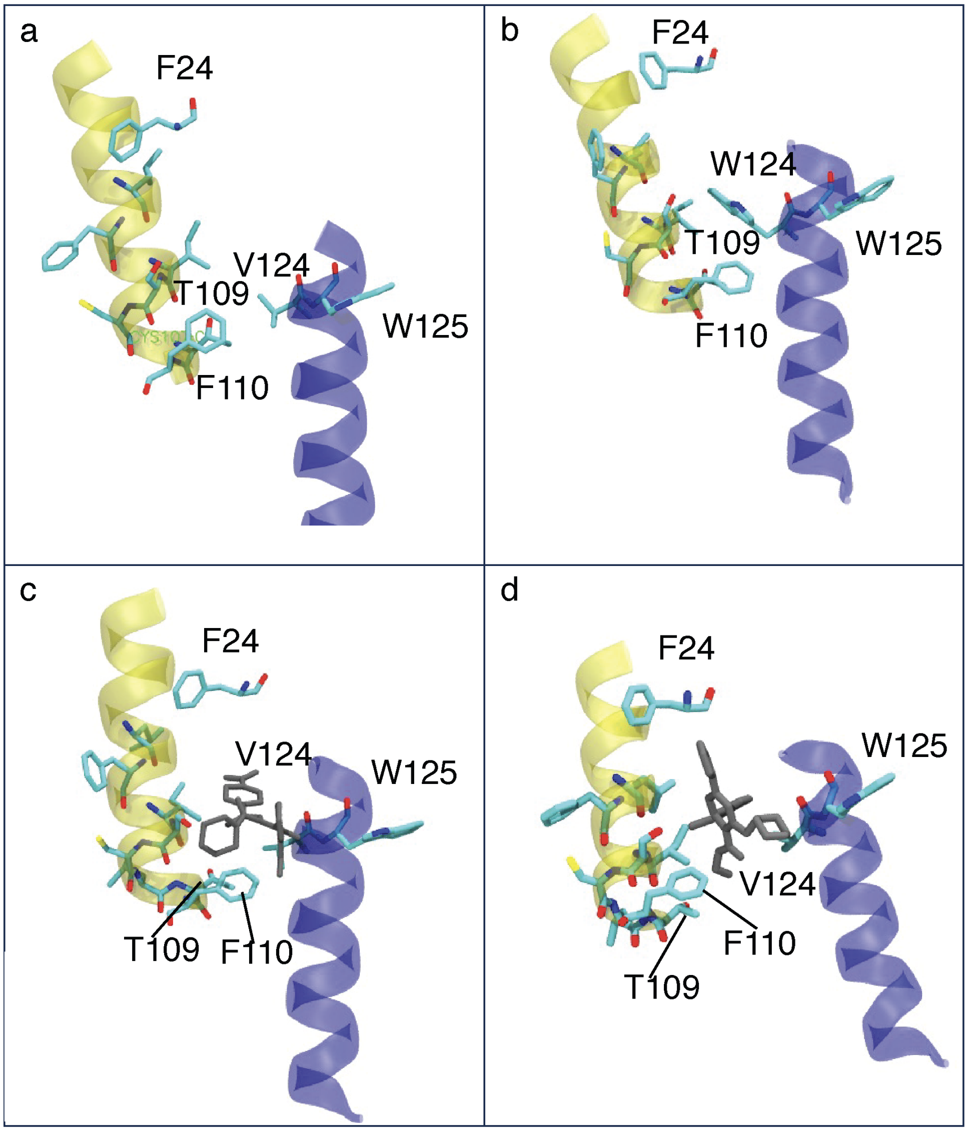


Figure S13: Comparison of the CAM binding site between ApoWT, Apo V124W, AT-130 bound and GLS4-bound states, selected snapshot from simulation is used for each case. In all figures, the secondary structure of C$\alpha$5 is shown in blue, and the secondary structure of B $\alpha$4b is shown in yellow. In addition, C residues V125 and V/W124, and B residues F24, L101, F103, I105, S107, L108, T109 and F110 and shown in licorice representation. These residues are shown to be importance by our residue-distance model when examining most important residue pairs. In all representations hydrogens are omitted for clarity. a) Structure of Apo system. b) Structure of V124W system. d) Structure of the AT130-bound system, AT130 is shown in grey. d) Structure of the GLS4-bound system, GLS4 is shown in grey.

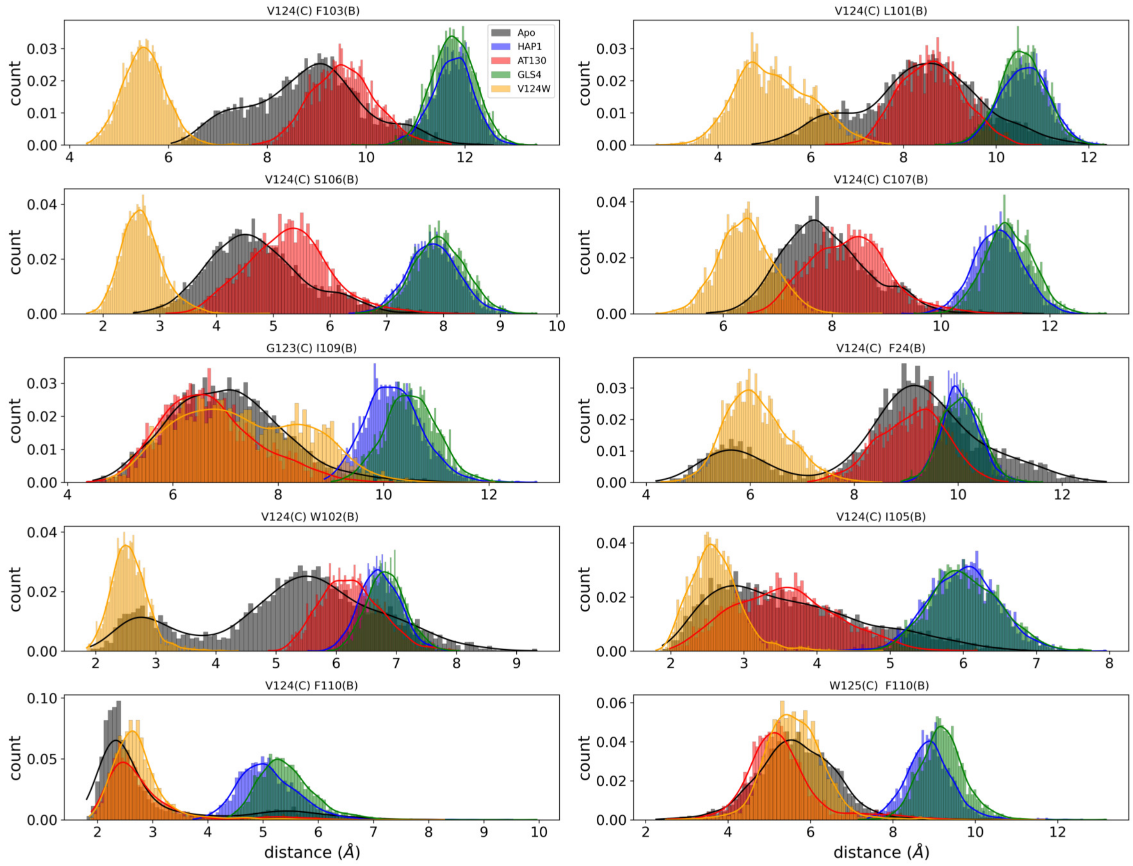


Figure S14: Distance distribution histograms for the top ten most important residue pairs in the residue-distance model.

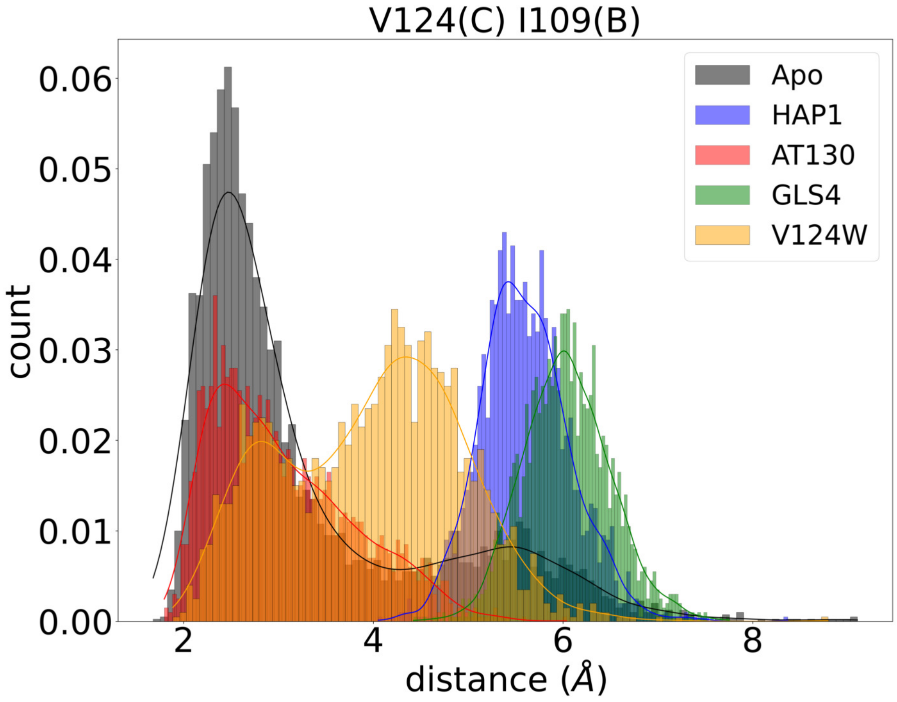


Figure S15: Distance distribution histogram for V124(C) and I109(B) residues.

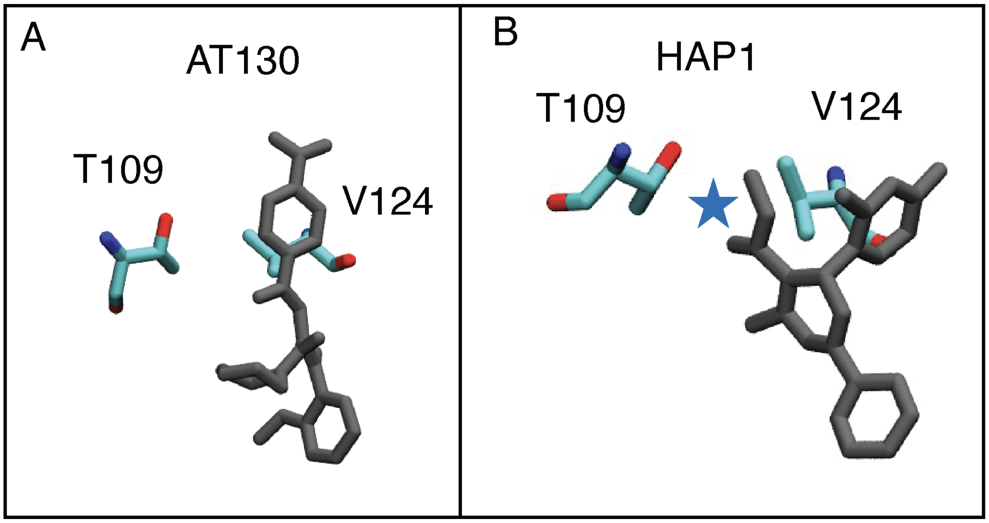


Figure S16: Comparison of binding between AT130 and GLS4 near residues V124(C) and I109(B). The compounds are shown in grey, and the ester group of GLS4 is marked with a blue star.
